# Eco-Evolutionary game dynamics with Three Strategies: how geometric analysis provides insights into stability

**DOI:** 10.64898/2025.12.26.696515

**Authors:** Manjyot Singh Bedi, Krzysztof Argasinski, Mark Broom

## Abstract

In this paper we extend a modeling framework that embeds ecological realism by examining how demographic factors determine payoffs, which are subsequently incorporated into evolutionary game models. In particular, it segregates births and deaths, as well as activities in addition to the between player interactions central to the classical theory. In the standard examples of evolutionary games so far considered such as the Hawk-Dove game, a two strategy population has been studied.

The above demographic models are extended from two strategies to three strategies, giving a thorough analysis of the mechanistic framework, which lays down the necessary conditions on the existence of steady states for a constrained system. We show that population trajectories are drawn toward the density manifold by logistic suppression, which in turn structures the interaction between frequency dynamics and fixed points. We also introduce a novel approach which helps derive stability from *geometric insights* through phase space elements (nullclines and their intersections) and streamlines calculations by decomposing gradients into the components that contribute to directional change, allowing dynamic and stability conditions to be more easily interpreted directly from the figures.

## 1 Introduction

Evolutionary game theory (EGT) is an important modelling methodology used to analyse the evolution of biological populations. Models are often elegant and explain apparently surprising features of nature [1–3]. Nevertheless, this simplicity comes at the price of realism and often the classical models were rather idealised. A series of papers [4–8], including by two of the authors of this work, has introduced the inclusion of explicit demographic parameters in an effort to make such models more plausible and biologically grounded. In this paper we build on that foundation.

EGT can trace its history back a long way to Düsing [9][10], but the classical theory is mostly attributed to Hamilton [2, 11] and especially Maynard Smith and Price [3], Maynard Smith [12]. In this classical approach [12–14], a well-mixed, unstructured and infinite population under natural selection is considered. Reproduction is clonal and there is no mutation; strategies are heritable phenotypic traits or behaviours and evolution occurs due to payoff functions incorporating an abstract measure of fitness. Maynard Smith was fully aware of the limitations of this approach, and the pros and cons are discussed in [12, 15–17].

The fitnesses considered, which can be further broken down into “cost” and “benefit” components, can be thought of as a vector of growth rates **r** and is described in undefined “units”. The simplest model involves a payoff matrix with *m* competing strategies, defined on the *m* − 1 dimensional simplex. Here, denoting *n*_*i*_ as the number of individuals playing the *i*th strategy, then *q*_*i*_ = *n*_*i*_/ ∑ *n*_*j*_ is the frequency of the *i*th strategy and *r*_*i*_(**q**) is its payoff function (where the population state **q** is the vector of strategy frequencies *q*_*i*_). The evolution of the population is commonly assumed to follow the replicator dynamics [14, 18–21]

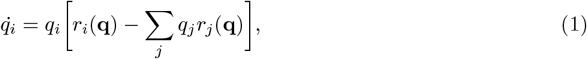

which is the most widely used evolutionary dynamics.

A more realistic game theoretic model of selection involving measurable demographic parameters was introduced in Argasinski and Broom [7], who provided a systematic approach to modelling “growth”. The excess over mean Malthusian growth or the use of “costs” and “benefits” is replaced by measurable outcomes of natural selection, births and deaths (similar approach was propsed in [22]). The base model is comprised of a frequency and a density equation for a single-population system, as dictated by the engagement of different kinds of fertility and mortality pressures. The demography is outlined by the chronology of events and the mechanistic interpretation [23] considers the probabilities of event outcomes as well as their rates of occurrence. When the chosen timescale is such that the interaction rate equals 1, the rates in the equation then correspond directly to the number of offspring and the probability of death resulting from an average interaction, expressed in the appropriate units (births and deaths per unit time). Integrating these demographic processes with simple matrix games and ecological parameters provides a unified framework for understanding the complex dynamics of the system.

In particular the model from [24], which is the basis of our work, was expressed in the following form

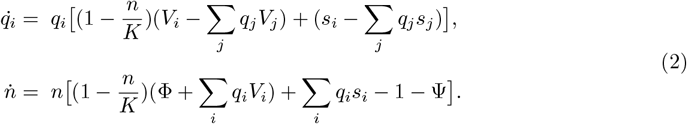

In addition to the population size *n* and strategy frequencies *q*_*i*_ already described, the model includes the following parameters:

The carrying capacity *K* describes the maximal population load bounded by the availability of some critical resource such as the number of nest sites [8, 25]; the reproduction and survival payoff functions *V*_*i*_ and *s*_*i*_ (these are both functions of **q** and can be represented as payoff matrices, as will be the case in this paper); background birth rate Φ and background death rate Ψ describing the impact of the factors other than the focal game. Whilst the theory leading to the demographic model (2) was developed in [24, 26], direct analysis was limited to a two strategy scenario based upon the Hawk-Dove game. Here we extend the analysis to consider the three-strategy case in generality, prove results and consider examples that show unique behaviours. In particular, the extension to three strategies led to us developing novel geometric methods to more easily explain the dynamic behaviour of the population.

## 2 A mechanistic framework to analyze three-strategy dynamics

Three strategy interactions are of significant interest and relate to different natural phenomena. The Rock-Paper-Scissors (RPS) game is the classic example of three-strategy cyclic competition which can be observed in nature, particularly in side-blotched lizards (Uta Stansburiana) and Coliform bacteria. Although it is simplistic in nature, its oscillatory impact coupled with evolution is quite complex [27, 28]. In three-strategy RPS dynamics, the interior coexistence equilibrium can be stable, unstable or neutrally stable, depending upon the payoff structure [29]. This leads respectively to convergence to stable coexistence, divergence toward boundary dominance or persistent cycling around the equilibrium.

Here we start with (2) where the *V*_*i*_ and *s*_*i*_ functions are in the form of 3 × 3 matrices of non-negative payoffs, representing the effects on fertility and mortality of interactions between the players.

### 2.1 The model

Our system is comprised of three frequency equations (which are effectively two, since ∑ *q*_*i*_=1) and a density equation. The dynamics are governed by two matrices: a fertility matrix (*A*), which highlights the reproductive contributions of each strategy and a survival matrix (*B*), which captures their respective survival rates for interaction events. These matrices, presented below, collectively establish the foundation for the model’s subsequent calculations:

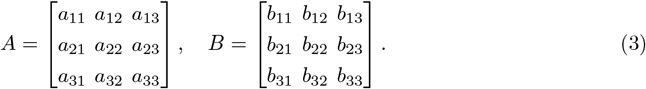

In the previous section, we outlined the differential equations for the evolution of the population for the general case. With *e*_*i*_ denoting the directional vector for each strategy, (2) becomes:

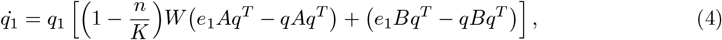

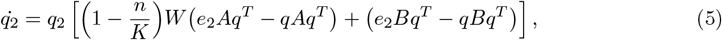

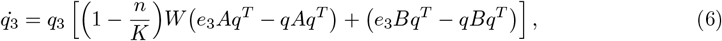

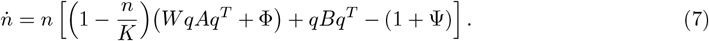

We note that in the above equations *W*, representing the fertility reward from an interaction, acts as a simple multiplier of the fertility matrix *A*, so that a suitable reparameterization of *A* would subsume it. But, given its natural interpretation, we will keep it as a distinct parameter.

Since the frequencies must sum to one, only two of them are independent and hence the frequency dynamics effectively have two degrees of freedom. Therefore, one frequency equation is redundant. However, later it will be useful in calculation of the nullcline of third strategy. A list of parameters and symbols (both, established so far and introduced later) are outlined in Table 1.

**Table 1:** Important symbols.

| Symbol | Description |
| --- | --- |
| **Section 2.1** |  |
| $A, B$ | Fertility and Survival payoff matrices |
| $q_i$ | Frequency of the $i^{\text{th}}$-strategy |
| $n$ | Population size |
| $K$ | Carrying capacity (maximum environmental load) |
| $W$ | Focal interaction fertility reward |
| $e_i$ | $i^{\text{th}}$-strategy directional vector |
| $\Phi$ | Background fertility rate |
| $\Psi$ | Background mortality rate |
| **Section 2.2** |  |
| $\alpha_i(q) := e_i A q^T$ | Fertility payoff of strategy $i$ against population state $q$ |
| $\beta_i(q) := e_i B q^T$ | Survival payoff of strategy $i$ against population state $q$ |
| $\bar{\alpha}(q) := \sum_i q_i \alpha_i(q)$ | Population weighted mean fertility payoff |
| $\bar{\beta}(q) := \sum_i q_i \beta_i(q)$ | Population weighted mean survival payoff |
| $c_1 = \Phi/W$ | Background-to-focal fertility ratio |
| $c_2 = 1 + \Psi$ | Effective baseline background mortality |
| $n^* = (1 - n/K)W$ | Density-dependent fertility multiplier |
| $\hat{n}_i$ | Population size on the $i^{\text{th}}$ frequency nullcline ($\dot{q}_i = 0$) |
| $\hat{n}$ | Population size on the density nullcline ($\dot{n} = 0$) |
| $\nu_i, \epsilon_i$ | Auxiliary notation encoding the pairwise interactions of strategies |
| **Section 2.3** |  |
| $\omega$ | Payoff proportionality parameter |
| $\tau_i$ | Parameter-dependent coefficients of the quadratic equilibrium condition in $q_2$ |
| **Section 3.2** |  |
| $\gamma$ | Tangent to frequency nullclines with gradients $\vec{q}_1$ and $\vec{q}_2$ |
| $\phi$ | Angle between any two chosen gradients |
| $\alpha_i^{(j)}$ | Angle between focal axis $i$ and adjacent nullcline, projected along orthogonal axis ($j$) |
| $\beta$ | Angle between gradient $\vec{n}$ and tangent $\gamma$ |
| **Section 3.3** |  |
| $\tilde{n}$ | Equilibrium point density |
| $\tilde{q}_i, \tilde{q}_i$ | Equilibrium point frequencies |
| $S_g(A)$ | Slope of the gradient A with respect to its own axis |
| $S_A^B$ | Slope of nullcline A along axis B |
| **Section 4** |  |
| $r_i$ | Focal payoff of the $i$th strategy consisting focal fertility and survival components |
| $\bar{q}$ | Average payoff of all strategies |
| $\rho$ | Background payoff consisting of background fertility and survival components |

### 2.2 The static interpretation

The original system consists of three frequency variables (*q*_1_, *q*_2_, *q*_3_), representing the relative magnitudes of each strategy, together with a density variable (*n*) denoting total population size. Population size is constrained by the logistic suppression coefficient, which ensures trajectories are drawn toward the population manifold. On this manifold, (*n*) can be expressed as a function of the frequencies, so the population is demographically stable and the evolutionary dynamics unfold in reduced dimensions. At equilibrium, the four-dimensional system (*q*_1_, *q*_2_, *q*_3_, *n*) therefore reduces to two-dimensions (*q*_1_, *q*_2_), with the density lying on a globally attracting manifold uniquely determined by the independent frequencies.

In this reduced setting, define for each strategy *i* ∈ {1, 2, 3}:

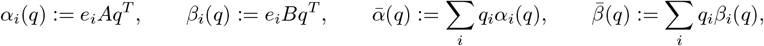

with *A, B* the fertility and survival matrices. Let *c*_1_ = Φ/*W*, *c*_2_ = 1 + Ψ and *n*^∗^ = (1 − *n*/*K*)*W*. Thus, the nullclines (curves in the phase plane where the rate of change of one variable is zero) are given as,

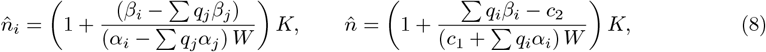

where 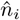 is the nullcline of *i*th strategy and 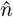 is the attracting density nullcline (also termed stationary density surface [30]). We see that the equilibrium conditions compress into a single set of equalities

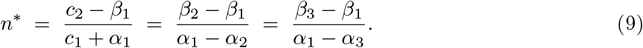

*For proof see Appendix A*

#### Lemma 1

(Background balance) *At equilibrium, the ecological constants* (*c*_1_, *c*_2_) *and the strategy payoffs satisfy the following background balance conditions:*

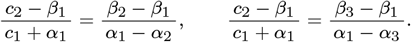

*In symbolic form these conditions reduce to two quadratic equations in the strategy frequencies* (*q*_1_, *q*_2_):

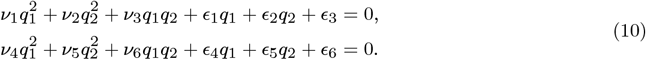

*Here the coefficients ν*_*i*_ *and ϵ*_*j*_ *denote auxiliary notation encoding the pairwise interactions of strategies (for notation, see Table A1*).

*For proof see Appendix A*

Now we obtain a clearer two-dimensional view of how the frequencies interact under the constraint of a stable population size, thereby defining the strategic composition of the system. Since the equilibrium conditions are expressed symbolically, they make transparent the fundamental influence of the fertility and survival matrices, allowing us to analyse the underlying game–theoretic structure in its most mechanistic form.

When we shift our focus from the density manifold to the the frequency nullclines (curves along which the growth rate of a particular strategy is zero) which span the full phase space, their intersection identifies strategy combinations that will remain stationary for fixed population sizes. Crucially, this intersection does not collapse to one equilibrium but rather it forms a continuous set of solutions, a line of strategy compositions. Each point from this set can be interpreted as a Nash equilibrium.

#### Theorem 1

(Line of Nash Equilibria) *Equilibrium in the frequency dynamics requires that all strategies yield identical effective payoffs, so that none has a selective advantage. This condition forces the ratios of payoff differences to align:*

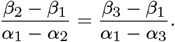

*Geometrically, the locus of points q* ∈ Δ^2^ *satisfying this equality is the zero set of a quadratic in* (*q*_1_, *q*_2_),

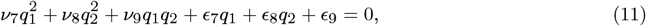

*with coefficients ν*_*i*_ *and ϵ*_*j*_ *determined by the structure of the fertility and survival matrices A and B (see Appendix A*).

*Every interior solution on this quadratic curve corresponds to a mixed Nash equilibrium, a strategic composition where no strategy can invade the others*.

*For proof see Appendix A*

#### Corollary 1

(Density independence) *The coefficients in* (11) *are determined entirely by the payoff matrices A and B. Consequently, the Nash line depends only on strategic interactions and is invariant under changes in population size or ecological parameters that enter solely through the influence of the suppression factor n*^∗^ *in* (9).

#### Corollary 2

(Eco-evolutionary equilibrium) *A frequency vector q on the Nash line constitutes a full ecoevolutionary equilibrium if and only if n*^∗^(*q*) *(obtained from any pairwise indifference condition) also satisfies the background balance equation* (9)

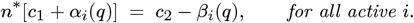

*Thus, the ecological constraint singles out discrete points on the continuous Nash line as genuine equilibria of the full system*.

### 2.3 A constrained system

A closed-form solution to the full symbolic system is not attainable. Such a solution, however, is highly desirable as it makes transparent how distinct classes of equilibria depend on the underlying parameters. To recover explicit expressions, we restrict attention to a subspace of the parameter space by imposing the additional condition in equation (12). This constraint is sufficiently mild to retain a rich and varied solution structure, yet strong enough to render the system analytically tractable. The construction of the constrained system centres on the two quadratics arising from the background balance conditions in equation (10). This choice is structural, as trajectories are confined to the density nullcline and the dynamics unfold in its vicinity. The third quadratic, in contrast, enforces the intersection of the frequency nullclines, which is purely strategic ensuring equal payoffs across strategies. It describes the geometry of equilibria along the Nash line, but does not necessarily play a role in pinning the dynamics to the density nullcline itself.

#### Definition 1

(Constraint Parameter *ω*) *We now define the parameter ω by imposing the condition*

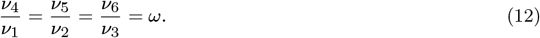

This restriction simplifies the system from (10), allowing explicit closed-form solutions for *q*_1_ and *q*_2_, capturing all possible mechanics observable for an interior fixed point and helps in simpler yet detailed calculations.

The fixed points of this system are conditional on the solution of the quadratic in *q*_2_, arising from the intersection of the two frequency nullclines under a fixed population size. Each coefficient (*τ*_*i*_) is a nonlinear combination of the system parameters and interaction terms. The equation is:

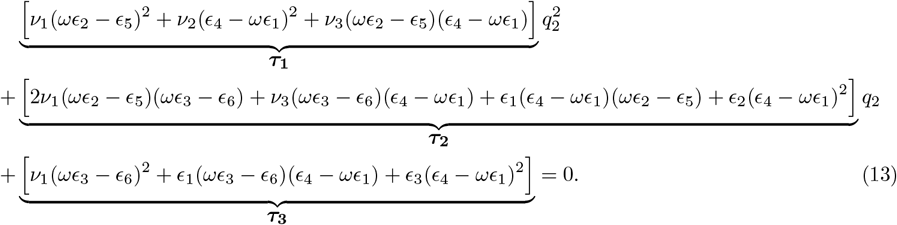

Upon determining the admissible root(s) for (*q*_2_), the corresponding values of (*q*_1_) are obtained from the following linear expression,

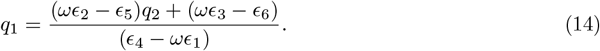

*For proof see Appendix B*

A special case of our approach arises when the differences in payoffs between the first and third strategies are proportional to the corresponding differences between the second and third strategies, across both payoff matrices. Specifically, when the relative magnitudes of strategies 1 and 2 are compared against strategy 3 using matrix *A*, we get:

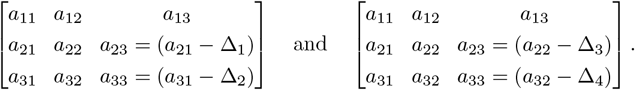

Assuming a constant proportionality *ω* implies the following chain of relationship:

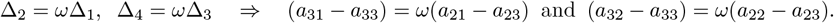

Similarly, we establish the proportionality *ω* for the second payoff matrix (*B*). Thus, our final unified condition predicated on the relative differences in payoffs between strategies, when measured against a common reference across both matrices, is:

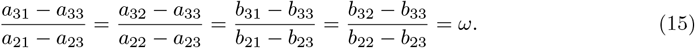

#### 2.3.1 Criteria for fixed points

Solutions to the system governed by equations (13) and (14) must be biologically relevant and to ensure real-valued solutions, the discriminant of the quadratic equation must be non-negative (i.e.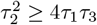). Therefore, feasibility within the frequency simplex is established via:

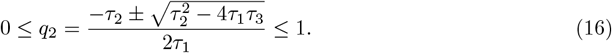

These criteria collectively define the admissible range of solutions for *q*_2_ and in turn the subset of parameter values that allow an equilibrium in the interior of the phase space (*q*_1_, *q*_2_, *q*_3_). Accordingly Table 2 summarizes the requirements for the system to yield valid fixed points under a stable population size:

**Table 2:**
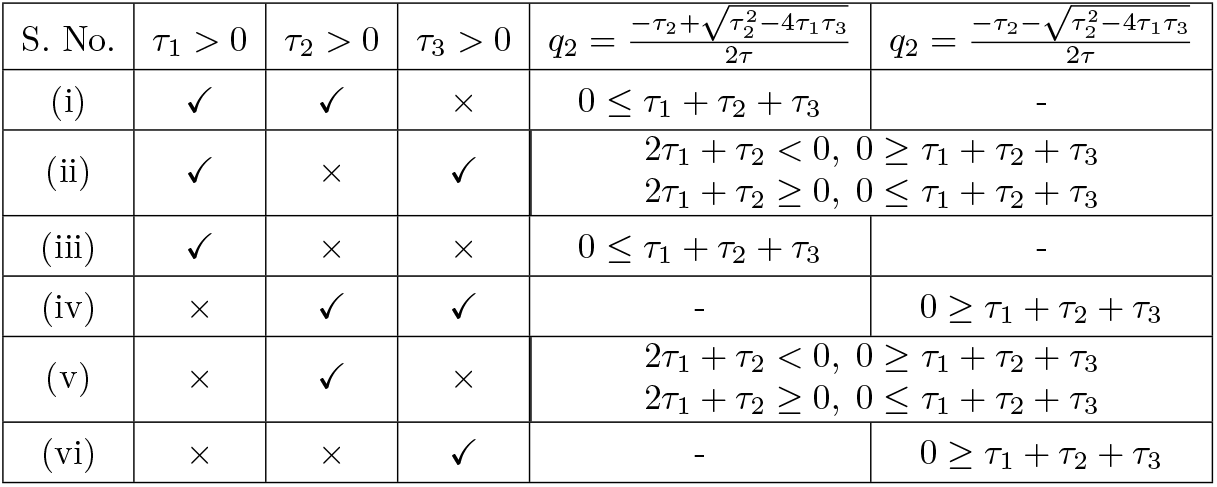
Conditions for existence of roots of *q*_2_.

### 2.4 Analysis of stability

The Jacobian matrix provides the local linear approximation of the system near its equilibrium. Its entries are partial derivatives of the dynamic variables with respect to one another and its eigenvalues determine stability: if all have negative real parts, the equilibrium is locally asymptotically stable; otherwise, the point is unstable or oscillatory. Thus it compactly encodes the interaction between strategy dynamics and population growth and provides the basis for stability analysis. A natural understanding of the stability conditions, centred on the attractor density surface, is achieved when the Jacobian follows a specific orientation of the phase space (*n, q*_2_, *q*_1_) which is opposite to the original.

From our replicator system (4), (5) and (7) we get,

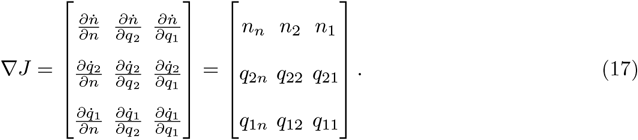

The Jacobian can be viewed as the transpose of the gradients of the right-hand sides of the equations. Each gradient is the normal vector to the nullcline of its corresponding equation. This geometric property will be central to our exposition in the following sections, where stability analysis is developed in terms of nullclines and their intersections. Subsequently, we obtain the characteristic equation as:

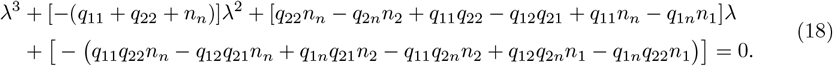

Stability of a system requires that all roots of its characteristic equation must have negative real parts. This condition can be assessed using the Routh-Hurwitz criterion (R-H conditions) [31, 32], which states that a 3 × 3 real matrix *J* is stable if and only if all of the following hold:

- Decaying Perturbations: Trace, *tr*(*J*) < 0, i.e.

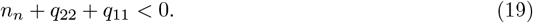

The trace corresponds to the sum of the eigenvalues of the system, which in turn determine the local behaviour of the system near an equilibrium point. A negative trace implies that trajectories converge to the equilibrium, ensuring stability. A positive trace indicates divergence and instability, while a zero trace suggests possible neutral stability, oscillations, or more complex dynamics that require further analysis.
- Steady Sub-System Interactions: Sum of principal minors, *a*_*t*_ > 0, i.e.

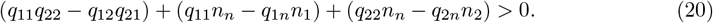

The principal minors capture pairwise interactions between state variables through the 2 × 2 determinants along the leading diagonal. A positive sum of principal minors indicates that the system is resistant to runaway growth, i.e. small perturbations remain bounded and the dynamics return to equilibrium.
- Partially Stable System: Determinant, *det*(*J*) < 0, i.e.

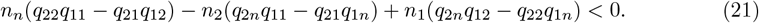

A positive determinant implies at least one unstable direction. Specifically at least one eigenvalue must be positive, implying saddle-point like behaviour in the phase space or an unstable node. A negative determinant is thus required for stability, though it is less conclusive, as it corresponds to an odd number of negative eigenvalues (one negative and two positive or all three negative), so the system may still be unstable and hence needs to be confirmed with the remaining conditions.
- Stabilizing local subsystems must resist destabilizing cyclic feedback loops: *tr*(*J*) ∗ *a*_*t*_ < *det*(*J*), i.e.

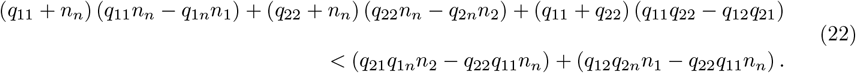

*For proof see Appendix C*

The expanded form of the final R–H condition separates stability into local and global contributions. The left-hand side, involving the sum of products of the principal minors and their respective traces, reflects local stabilizing interactions. This hints at self-regulation and dissipation between variable pairs. The right-hand side, built from products of off-diagonal elements along cyclic triangular paths, represents global destabilizing feedback loops. These products are termed the (signed) weights of circuits, which are defined as sets of Jacobian elements whose indices form a cyclic permutation [33]. Stability is achieved when the cumulative strength of local stabilizing interactions outweigh the destabilizing potential of global cyclic feedbacks.

## 3 The geometry of stability

Since the Jacobian is a transposed matrix of gradients, stability can be understood not only through algebraic criteria, but also through a geometric lens that highlights the spatial relationships underlying the system. At the heart of this perspective lie the systemic gradients given by **q**_**1**_ = (*q*_11_, *q*_12_, *q*_1*n*_), **q**_**2**_ = (*q*_21_, *q*_22_, *q*_2*n*_) and **n** = (*n*_1_, *n*_2_, *n*_*n*_).

Each of these vectors serves as a normal to the surface defined by a corresponding nullcline, so that the geometry of their mutual orientation encodes how the system evolves near equilibrium. Dot products, cross products and projections provide the algebraic tools for capturing these orientations, translating the original stability conditions into relationships among gradients, or alternatively, nullclines.

Our approach considers the system in its full dimensional setting. From there, dimensional reduction reveals how stability manifests step by step, making the role of each geometric component explicit. This layered view uncovers the structural interplay of gradients and offers a complementary intuition for why and how stability is maintained.

### 3.1 Stability from cross and dot products of gradients

- **Three-dimensional condition**: Nullclines define surfaces where the rate of change of each respective variable is zero. When two such nullclines intersect, the intersection forms a line along which the equilibria of the two-dimensional subsystem (with the third variable fixed), consisting of those two equations, reside. In our case, we can choose the frequency nullclines. Then the intersection constitutes the set of Nash equilibria. The stability of these equilibria depends upon the local behaviour of the system near the intersection, determined by the orientations and magnitudes of the gradients. In this geometric view, the cross product from (24) when dotted with the **n** gradient yields a scalar quantity reflecting this volume and its directional consistency with system stability as follows:

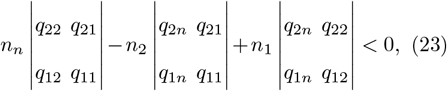

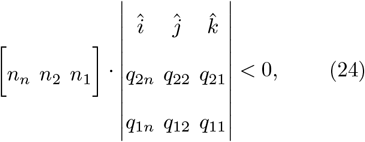

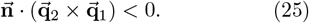

The cross product of gradients **q**_1_ and **q**_2_ in (25) refines our understanding of the system’s stability. Geometrically, the angle between these two gradient vectors determines the *direction of the resulting cross product vector*, which is orthogonal to the plane defined by **q**_1_ and **q**_2_. At a nullcline intersection, this cross product indicates whether the combined gradient effect points *upwards or downwards along the line of intersection (constituting the Nash equilibria in our case)*. Since the population-size gradient **n** typically points downward relative to the density nullcline, stability depends upon alignment: if the frequency gradients’ cross product points to the opposite side, the Jacobian determinant is negative and the equilibrium is stable. Conversely, if the cross product also points downwards, it aligns with the direction of the **n** gradient, potentially leading to instability.
- **Two-dimensional condition**: Each principal minor of the Jacobian matrix, from (26), has a natural geometric interpretation as a projection onto the surface spanned by a particular pair of state variables, as can be seen from (27). Then, for example, the phase subspace spanned by the **q**_1_ and **q**_2_ axes can be visualised as the *floor* or a reference plane, with each gradient representing directional changes along that floor. The cross product of the projections of the two gradients yields a vector whose magnitude represents the area of the parallelogram formed by the two gradients and whose direction (denoted by the unit vector) is orthogonal to the plane spanned by them. The area of the parallelogram equals the area of the square spanned by one of the gradients and the height of the parallelogram. Thus, dividing this magnitude by the norm of one of the gradients yields the height of the parallelogram, which corresponds to the projection of the second vector onto the direction perpendicular to the first projected gradient. The sign of the cross product indicates on which side of the nullcline surface the projection lies, distinguishing whether the nullcline intersection acts locally as an attractor or a repeller. When the cross product changes sign, it indicates that the gradients have become collinear, meaning the nullclines coincide entirely at that point and stability properties switch. Thus, each principal minor not only contains algebraic conditions necessary for stability, but also reflects the geometric interplay of the gradients’ orientations, the projection directions and the resulting dynamical behaviour near equilibria, showing how the system’s state space is structured and how perturbations evolve within it. Thus, the principal minor condition expressed in terms of determinants and corresponding cross products will have the form (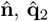 and 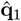 are respective axes unit vectors):

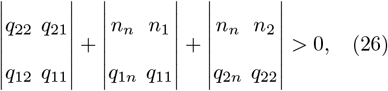

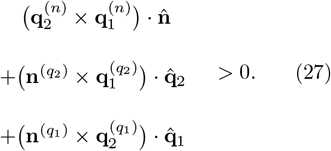
- **One-dimensional condition**: The trace gives the divergence of the vector field at that point. It is the one-dimensional rate of total expansion along coordinate axes. The trace element for each variable in (28) represents its contribution to its own change. So it can be viewed as projections of entire transformations onto the main coordinate axes and adding up the line segments. If each axis shortens, the trace is negative and hence the system experiences damping.

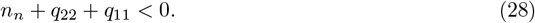
- **Cross-dimensional condition**: Up until now, we viewed each of the separate components of the stability conditions as vectors, building on their remote alignments in the phase space. The final condition from (22), involves their combined interactions as follows,

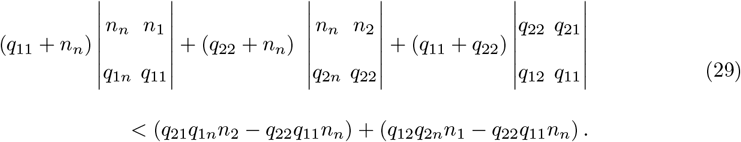

### 3.2 Stability via lengths of the gradients and angles between them

At any intersection, two complementary angles exist, so a consistent orientation must be chosen to ensure compatibility with the corresponding angle between gradients. In the three-dimensional space, the Nash line is defined as the intersection of two frequency nullclines with gradients 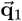 and 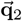, whose tangent is,

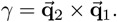

Note that the floor projection of this vector, 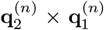, determines the orientation of the Nash line along the vertical *n* axis co-directional with *γ*.

Then the three dimensional condition can be denoted as:

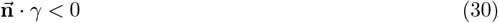

Where 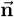 denotes the normal to the density nullcline.

Assume that *ϕ* denotes the angle between any two gradients and that when they are projected on one of the planes, the superscript of the orthogonal axis is added as can be seen in Figure 6. The remaining angles on the projected plane are denoted by *α*_*i*_, where *i* represents the focal axis which completes the angle. In addition *β* is the angle between the **ñ** gradient and the vector tangent to the Nash line resulting from the cross product 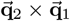. Then the stability conditions can be presented in terms of the angles between nullclines as:

**Fig. 1:**
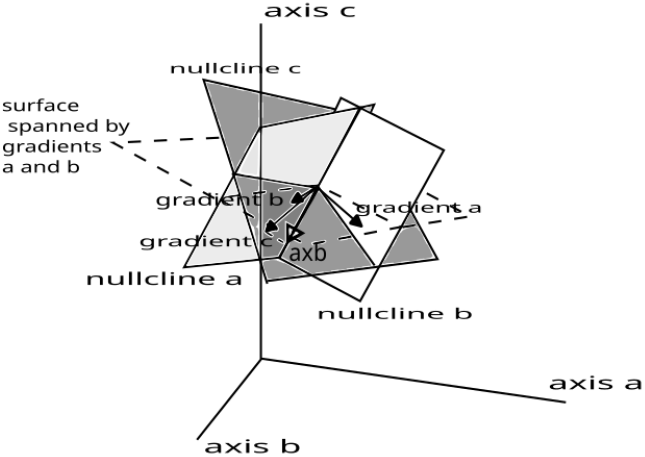
Geometric interpretation of the determinant. The cross product of two gradients generates a vector parallel to the intersection of their nullcline surfaces. The determinant is negative when the third gradient lies on the opposite side of the surface spanned by the first two gradients relative to the side indicated by their cross product. Therefore in the situation presented in the figure, we have an unstable intersection.

**Fig. 2:**
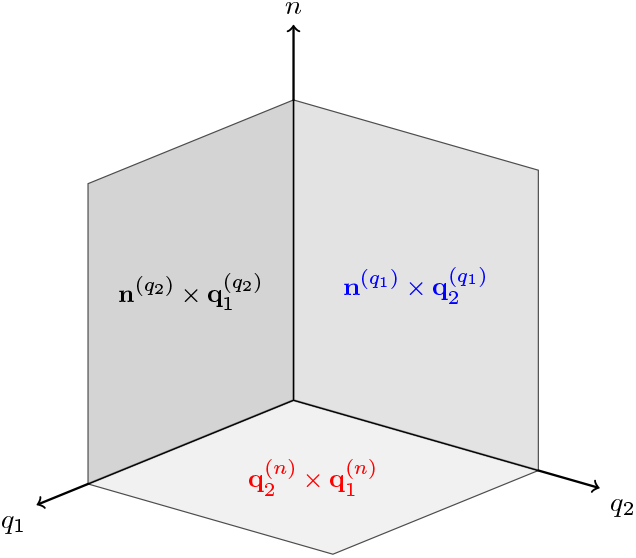
Gradient representations within the phase portrait. Each paired cross product corresponds to a principal minor, representing the “floor” and the “walls” of the system’s geometry. These cross products define the two-dimensional planes onto which the system’s dynamics are projected, while the third (described by bracketed superscript) is orthogonal to these planes.

**Fig. 3:**
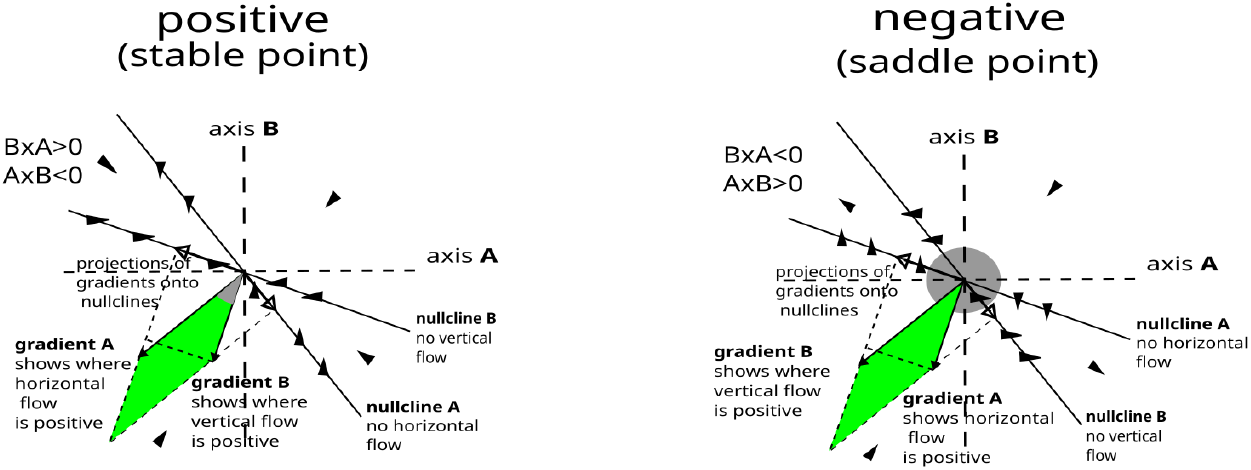
Geometric interpretation of the principal minor as the cross-product of the 2-dimensional projections of the gradients. The sign of the expression is determined by the direction of the projection of the opposing gradient onto the nullcline perpendicular to the focal gradient. This projection indicates where the flow (horizontal or vertical) induced by the opposite gradient is positive. Note that the clockwise angle between the gradients (given the chosen orientation (*n, q*_2_, *q*_1_)) equals the clockwise angle between their corresponding nullclines, allowing inference of their relative positions (which is higher or lower). The only difference between the two plots is the order of gradients and nullclines.

**Fig. 4:**
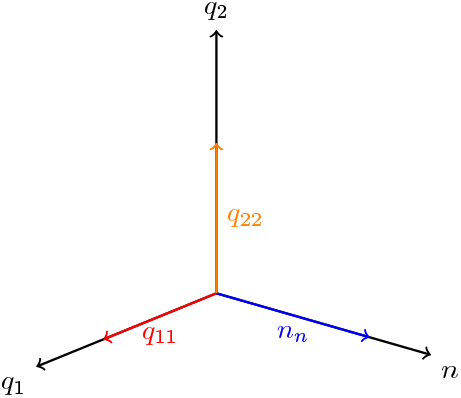
The trace components are shown as axis-aligned self-interactions. These self-interactions determine the local growth or decay rates along their respective directions within the phase space.

**Fig. 5:**
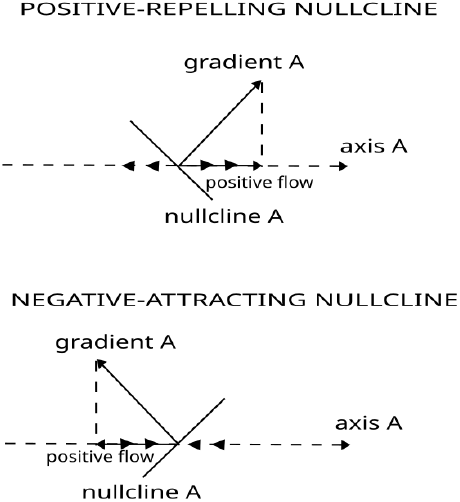
Geometric interpretation of the trace. The figure explains how the position of the gradient of the right-hand side of the equation describing the flow along a particular axis, relative to the focal axis, determines stability or instability of the respective nullcline.

**Fig. 6:**
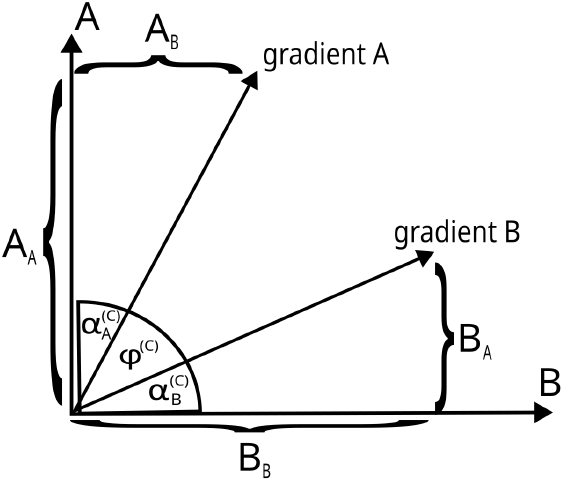
Angles corresponding to the projection along axis *C*, orthogonal to the plane defined by *A* and *B*.

**Fig. 7:**
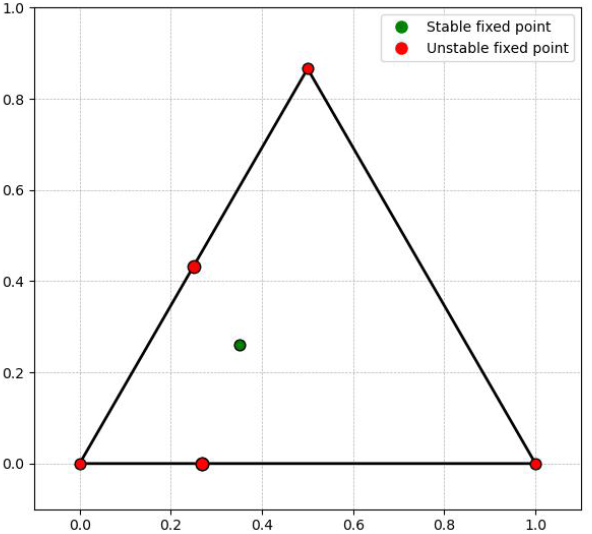
Example 1: Complete set of fixed points of the system, including both stable and unstable equilibria.

- **Three-dimensional condition:** Based on the angle between *n*-gradient and the vector tangent to Nash line, we have

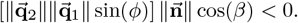
- **Two-dimensional condition:** Comprises the projections on the *floor* and *walls* as follows,

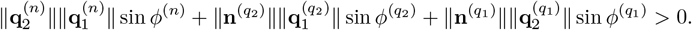
- **One-dimensional condition:** Remains unchanged as it is an algebraic result,

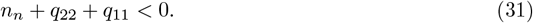
- **Cross-dimensional condition:** In terms of the angles and projections of gradients, (22) is presented as,

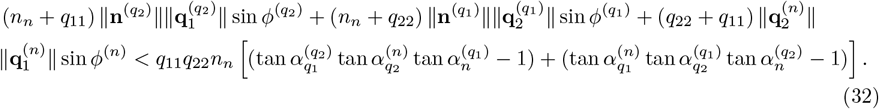

#### Remark 1

For the case with attracting nullclines (implying negative trace elements) then if, on every plane, the angle *ϕ*^(*C*)^ between the gradients *A* and *B* is sufficiently large so that both 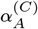 and 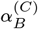 are less than 45^°^ (equivalently, 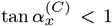, see Fig. 6), then the right-hand side of equation (32) is strictly positive, whereas the left-hand side is strictly negative. Consequently, the inequality holds trivially in such cases.

### 3.3 Stability through nullcline intersections and slopes

Assume that the Nash line, parametrized by the frequency as a function of the densities 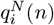, intersects the density nullcline at *ñ*. As the Nash line points upwards according to the direction of the cross product of the frequency gradients, it should locally lie above the density nullcline to achieve stability.

- **Three-dimensional condition:** This condition is satisfied when there exists an ϵ > 0 such that, for every *z* ∈ (0, *ϵ*),

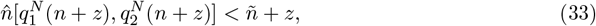

where *z* = 0 corresponds to the equilibrium intersection point (*ñ*). Subtracting 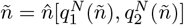 from both sides and dividing by *z*, the limiting form as *z* → 0^+^ yields:

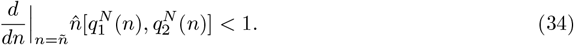

We note that it is not necessary to compute this derivative explicitly. Inequality (33) provides a practical alternative, avoiding the complexity associated with directly evaluating (34).

From Figure 6, the slope of the projection of the gradient *A* (on the plane spanned by axes *A* and *B*), with respect to its own axis is

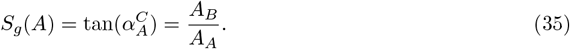

The corresponding slope of the nullcline along the opposite axis, denoted by 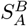 (where the subscript specifies the variable described by the nullcline and the superscript without brackets denotes the axis of measurement) is then:

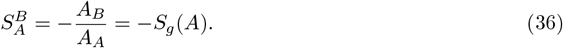

Similarly, the slope of the nullcline along the focal axis is

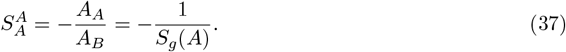

Then the stability conditions expressed in terms of the nullcline slopes will be:

- **Two-dimensional condition:** Using the definition of nullcline slopes, we have

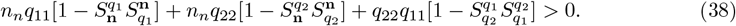
- **One-dimensional condition:** This condition remains algebraic and unchanged,

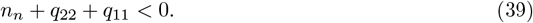
- **Cross-dimensional condition:** Based on the above understanding, (22) can be written in terms of the slopes of the nullclines, which for *q*_11_*q*_22_*n*_*n*_ < 0 (*q*_11_*q*_22_*n*_*n*_ > 0) can be presented as:

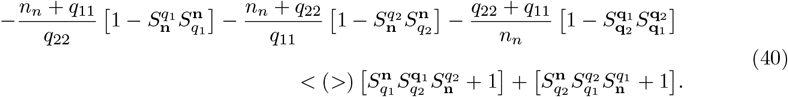

*For proof see Appendix D*

**Remark 2** For the case with attracting nullclines (implying negative trace elements) and the slopes of their respective projections are positive and less than 1, all conditions except the three-dimensional condition are satisfied only if the corresponding trace coefficients have signs consistent with the inequalities.

In more complex cases, however, the nullcline method can be useful for simplifying the calculations. Note that the slope products from the three bracketed terms of form 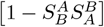 from (40) can, respectively, be presented as the ratios of the slopes along the chosen axis, leading to:

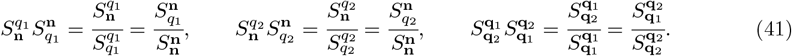

When both slopes have the same sign, for the corresponding bracketed term to be positive, the slope of the opposite variable’s nullcline should be smaller than the focal variable’s nullcline. When the slopes have different signs the bracketed term is positive trivially. Thus the nullcline slopes can be expressed with respect to the same focal axis, which yields

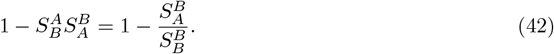

A possible situation is when the nullcline of the focal-axis dynamics *B* forms a straight vertical line. Thus, it cannot be regarded as a function of the variable *b*. However, in this case 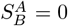, and the above bracketed term reduces to 1. In effect, the whole stability condition reduces to the product of the trace elements. The same occurs for the underlying cross product, since the vertical coordinate of one of the gradients is zero. The second extreme case is when the slope of the nullcline of variable **A** along the axis **A** is zero, which implies that the respective trace element is zero and the reciprocal nullcline slope in (42) is infinite. In this case, the slope condition cannot be evaluated. However, we can switch to the cross-product form, which collapses to the product of the elements of the second diagonal, since the trace contains zero.

In the following, a “wall” or “floor” projection of the nullclines refers to their values for a fixed parameter describing the coordinate along the axis orthogonal to the plane. Recall that we denoted nullclines as the manifolds 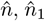 and 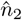 over the *q*_1_,*q*_2_ plane. Then, for example for the *n,q*_1_ plane, we can denote the two-dimensional sections of the nullclines 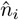 and 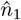 as either 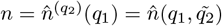 or 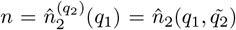. Similarly, for the “floor” condition, we can derive such sections from equations 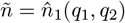 or 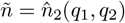 and denote them as 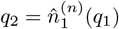 or 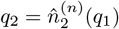. Let us focus on the “floor” case as an example. From the first-order Taylor expansion,

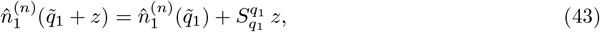

so the slope of the “floor” action of the nullcline of strategy 1, along the axis *q*_1_, can be approximated by

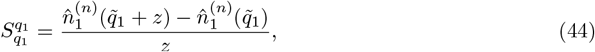

for sufficiently small *z*. This can be regarded as an “inverse Taylor approximation”, where we approximate an unknown slope from the values of a known function. In effect, we obtain

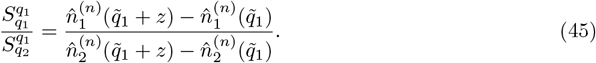

The other slope ratios can be computed in a similar way. The only complication arises in the final R-H condition (40) involving products of three slopes. For example,

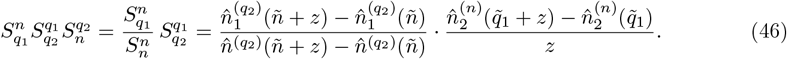

Thus, we can avoid complex derivative calculations and reduce the stability conditions to algebraic inequalities, which can be numerically evaluated over the parameter space when needed.

## 4 Gradient directional decomposition

Previous sections helped us to infer stability from nullcline geometry, leading to a substantial simplification of the required calculations. This approach allows us to determine *where* in parameter space the focal point is stable. However, we can go further by asking *why* stability arises for these parameter values by examining the underlying algebraic relationships that link the model parameters. In this section, we propose a methodology to isolate the necessary gradients into their respective components. The key idea is that gradients often contain information that is redundant or does not contribute to the actual direction of change in the system. By decomposing them into meaningful parts, we can focus only on the components that drive the dynamics. Thus *Gradient directional decomposition (GDD)* reduces computational complexity and highlights the essential mechanisms of evolution within the system.

GDD can, in principle, be applied to any system, but it is most effective when the vectors are aligned in a specific way. Mathematically, the cross product of two non-zero vectors is zero when they are parallel or anti-parallel, while the dot product is zero when the vectors are orthogonal. These geometric facts imply that the orientation in which the system evolves determines whether certain gradient contributions are active or redundant.

To make the notation precise, for each strategy *i*, we define the scalar *focal payoff*,

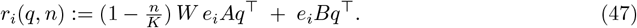

The background payoff appearing in the density equation is

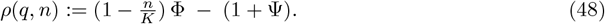

The population-level average payoff is the *q*-weighted mean of the focal payoffs,

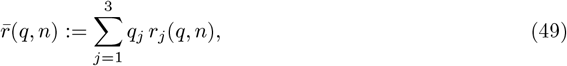

which, using the explicit form of *r*_*j*_, becomes

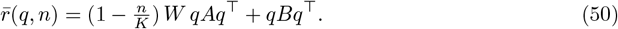

In our replicator system, the direction of change is governed by the relative movement with respect to the average 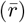. Intuitively, it can be written as:

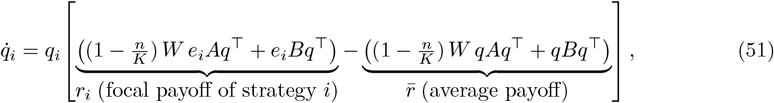

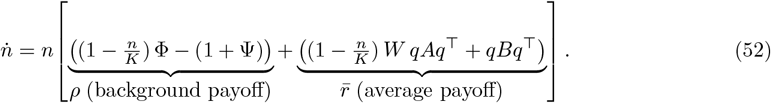

We denote by

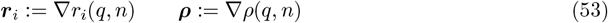

the corresponding *gradient vectors* of the focal and background payoffs. Superscripts such as 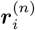 or 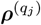, which appear below, denote projections of the full gradients onto the associated coordinate subspaces (e.g. the 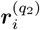 describes the projection onto the (*n, q*_1_)-plane along the *q*_2_ axis). Since gradient decomposition inherently simplifies each structural term, we do not need to apply it to the fourth R–H condition. It follows directly from the standard inequality *tr*(*J*) ∗ *a*_*t*_ < *det*(*J*) which can be evaluated most directly using the results obtained as follows:

- **Three-dimensional condition:** By applying GDD to the determinant of the Jacobian, specifically to (25), the analysis reduces from an inefficient expansion of mixed terms to a tractable form involving only the focal payoffs of strategies and their background contributions. Algebraically, the determinant decomposes into two interpretable parts - the triple product of the focal payoff vectors and the projection of the background payoff through the cross products of these vectors, as follows:

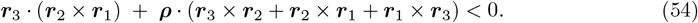

*For proof see Appendix E.1* This dual perspective shows how the geometry of alignment is directly tied to stability and can also be used to make the calculations more efficient by avoiding a full symbolic expansion of the Jacobian.
- **Two-dimensional condition:** Geometrically, the sum of principal minors can be seen as a balance between orientation volumes defined by payoff interactions and their projections along the strategy frequency directions. Algebraically, the gradient splitting framework when applied to (26) makes explicit how the stability condition decomposes into interpretable parts: focal payoff interactions, weighted by their contribution to the average and background-payoff corrections that adjust these interactions through cross products. The weights (*q*_*i*_) represent the relative contribution of each focal payoff strategy to the population average payoff 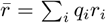, allowing the expression to be written in terms of weighted cross products of payoff vectors as follows:

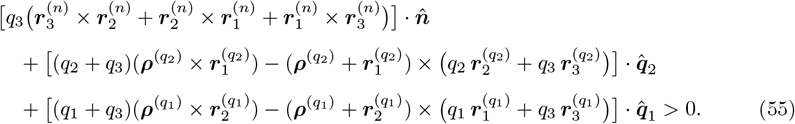

*For proof see Appendix E.2*
- **One-dimensional condition:** The trace is not a gradient, but rather a linear expression that captures the divergence of the vector field along each axis. In essence, it quantifies the net outflow rate at a point, rather than the slope. Although it is not an ideal candidate for gradient splitting, the resulting expression highlights that the fertility contribution to divergence is constant in frequency space but scaled by density dependence, while the survival contribution remains constant and independent of *n*, as can be seen in (56)

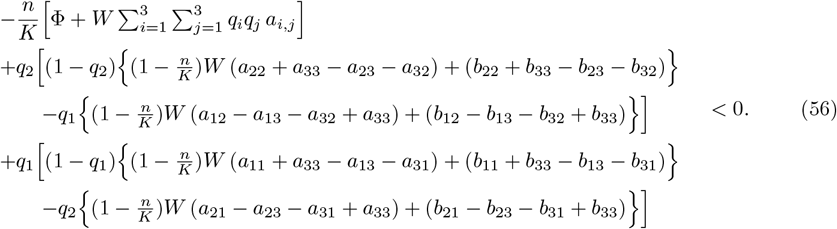

*For proof see Appendix E.3*

We have thus introduced three complementary propositions to help understand three-strategy dynamics. First, by imposing an exogenous constraint, we gained analytical tractability while retaining flexibility to observe the full range of dynamic behaviours around an internal fixed point. Secondly, we developed a geometric method that allows stability assessment directly through the phase space elements, offering insight without the need for explicit solutions. Thirdly, we proposed a calculation technique that systematically removes redundant or non-contributing components of the dynamic vectors, thereby isolating the active drivers of the system.

These propositions are novel in that they function both as stand-alone tools and as an integrated framework. The constraint sharpens focus on internal equilibria, the geometric method reveals system behaviour via nullcline interactions and gradient decomposition highlights the effective contributions to dynamics. Together, they provide a flexible and coherent approach for analytical and geometric study, underscoring the originality and practical utility of our work.

## 5 Numerical examples

We now apply our model to some concrete examples in order to illustrate the methodology for a stable interior solution and an unstable one. Then we verify the existence of fixed points highlighting boundary and vertex steady states. We consider three examples; in all of these we have the following four parameter values: *K* = 1000, *W* = 3, Φ = 1 and Ψ = 0. The other parameters are given in the payoff matrices *A* and *B* near at start of each example.

### 1. Stable interior solution

The trajectories that start from the interior of the simplex, where all three subpopulations exist initially, converge to a single interior fixed point as can be seen in cell 23 in Appendix F. An interesting observation lies in how the dynamics are offset by the use of the constraint parameter *ω*.

The payoff matrices are :

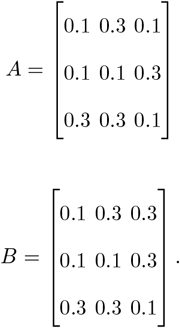

- **Constrained system calculations**

The above matrices represent the interactions of the three strategies, with each row and column encoding payoffs to the respective strategy. On substitution into the framework developed in Section 2.2, we obtain the following transformed system:

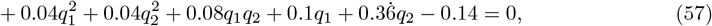

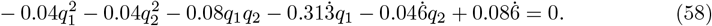

This system satisfies the condition from (12), since

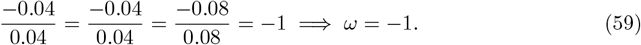

Consequently, the corresponding relations from (13) and (14) are,

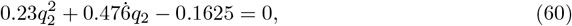

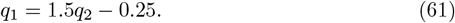

Table 3 verifies the admissibility of the roots and confirms their consistency with the theoretical criteria established in Table 2.

**Table 3:**
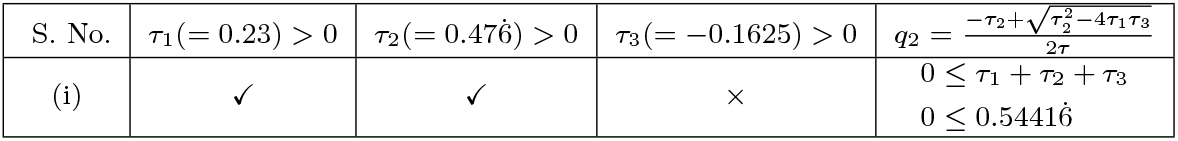
Corroborated conditions for roots of *q*_2_ for example 1.

Solving explicitly, the equilibrium values are *q*_1_ = 0.2, *q*_2_ = 0.3, *q*_3_ = 0.5, *n* = 5000. So far, we have confirmed the algebraic results for the constrained system. We now turn to its phase space. Figure 8 shows the time evolution of the three strategies converging toward a stable distribution, while Figure 9 presents a heatmap of the phase portrait, highlighting both direction and speed of convergence.

**Fig. 8:**
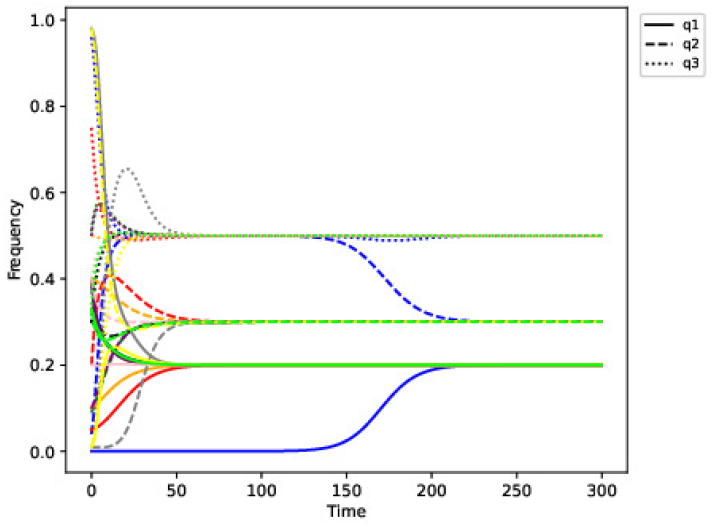
Example 1: Evolution of strategy frequencies over time. The plot shows the temporal dynamics of three strategies represented by a solid line, dashed line and dotted line, respectively. As time progresses, the system converges to a long-term stable frequency distribution.

**Fig. 9:**
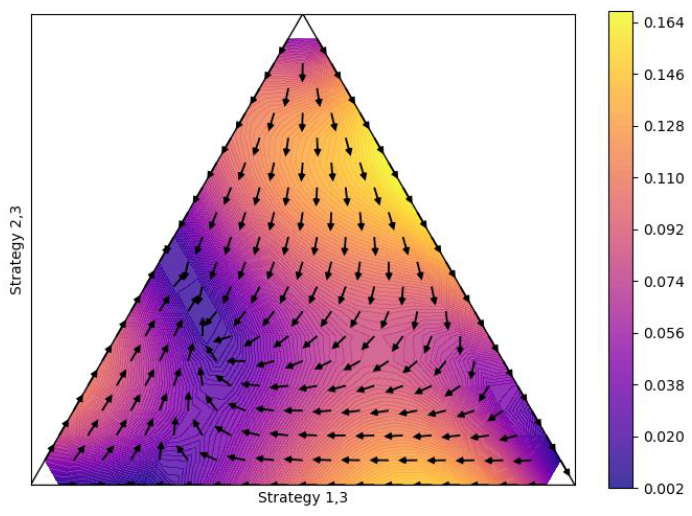
Example 1: Heatmap of the phase portrait illustrating the direction and speed of convergence. Arrows indicate the flow of the system, while colour intensity encodes stability - darker regions denote stable areas with stronger convergence and lighter regions indicate instability or slower dynamics.

Figures 10 (3d view) and 11 (overhead view) show the phase portraits with example trajectories. We see regions of slow dynamics, which appear to occur along the flow lines connecting the boundary saddles to the interior equilibrium. These trajectories resemble heteroclinic connections and the reduced speed along them is consistent with the system’s approach to saddle-type rest points. The heteroclinic orbits are related to the dynamics of the invasion of the rare mutant. This situation is illustrated by the blue trajectory on Figure 8, where the initial frequency of the first strategy is very low. Initially, the dynamics rapidly converge to the close vicinity of the unstable rest point followed by the slow convergence along the heteroclinic orbit to the interior stable equilibrium.

**Fig. 10:**
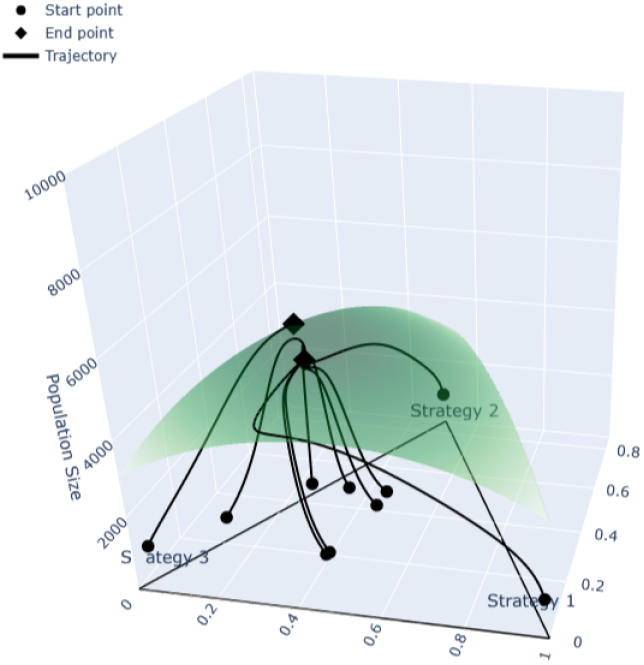
Example 1: 3D phase portrait of example trajectories: interior orbits approach the density nullcline and then converge to the stable equilibrium. The boundary trajectory instead converges to a boundary unstable equilibrium (stable in the reduced two-strategy system). A rare mutant with the third strategy drives the state along the heteroclinic connection toward the interior stable equilibrium.

**Fig. 11:**
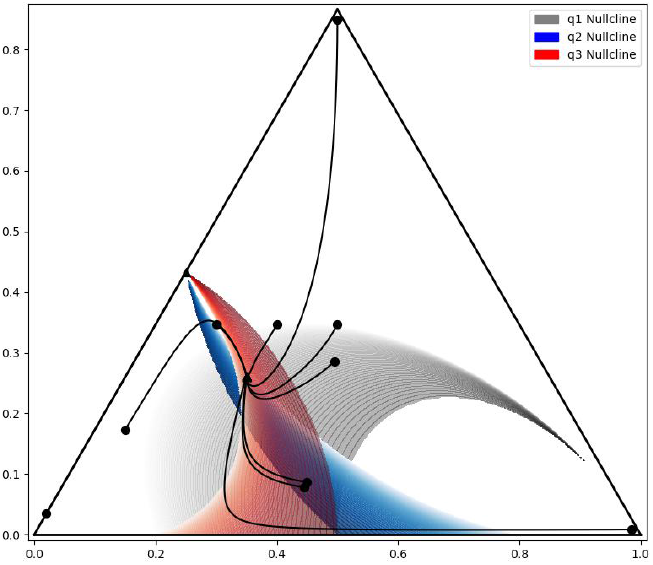
Example 1: Overhead view of trajectories superimposed on nullclines. The convergence of trajectories toward the intersection highlights the location of a stable fixed point, where the system’s dynamics settle into equilibrium.

- **Geometric Analysis**

The natural progression in our framework is to examine the geometry of nullclines and equilibria. Figure 12 shows the 3D nullcline intersection along with the Nash line, while Figure 13 provides isolated views of each nullcline. Together, these highlight how the equilibrium is determined by the joint structure of the system. Figures 14, 15 and 16 show the geometry of the respective principal minor conditions and demonstrate that the system is attracting in every plane.

**Fig. 12:**
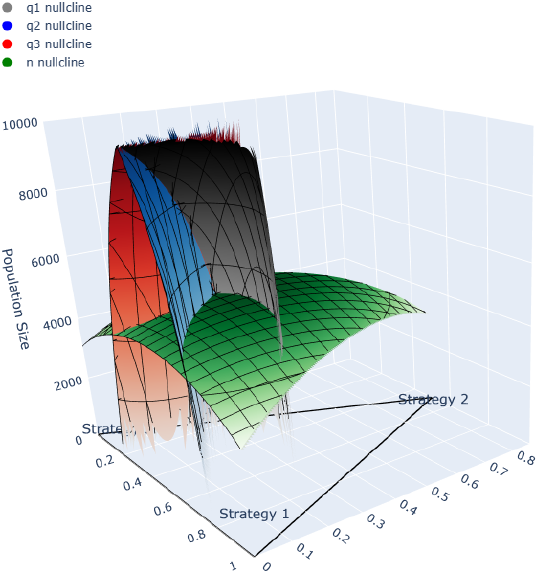
Example 1: 3D nullcline intersection. The intersection of the three frequency nullclines indicates that for each population size, there exists a corresponding steady frequency distribution. Together they form a continuous equilibrium line (a Nash line) across population levels.

**Fig. 13:**
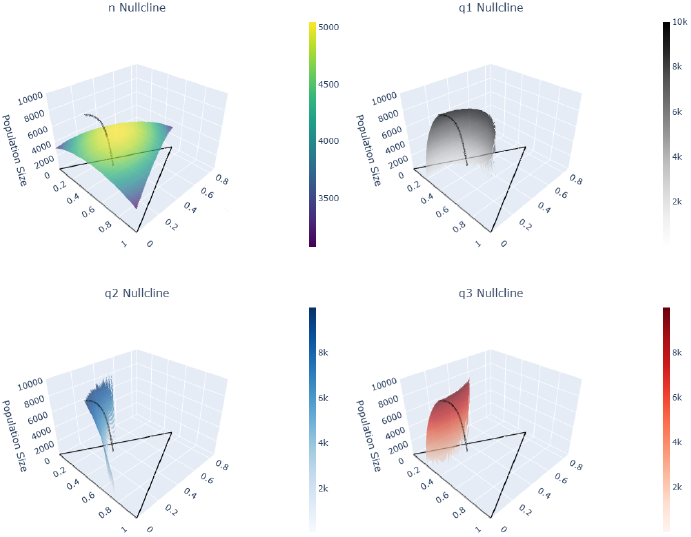
Example 1: 3D illustration of individual nullclines, with line of Nash Equilibria(from top left, clockwise): Population nullcline, *q*_1_ nullcline, *q*_3_ nullcline, *q*_2_ nullcline. Each subplot isolates a specific nullcline to highlight its structure and behaviour within the phase space.

**Fig. 14:**
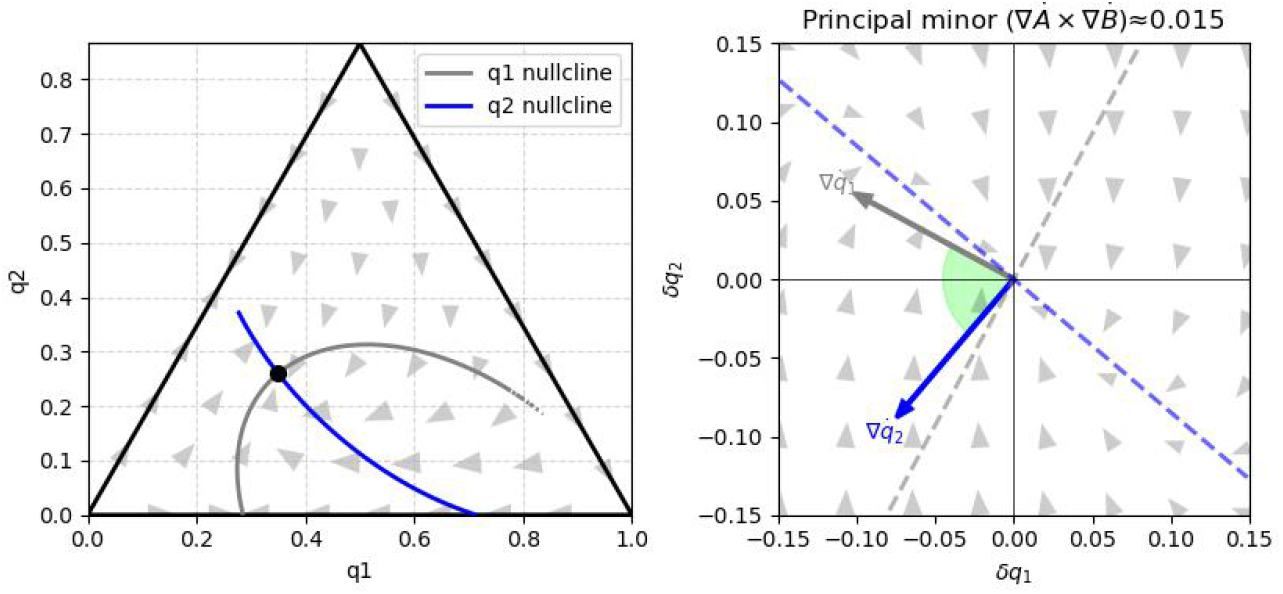
Example 1: “Floor” section for the *q*_1_, *q*_2_ principal minor. Left: flow and nullclines. Right: zoomed intersection in Cartesian coordinates with gradient angle, indicating stability in this plane.

**Fig. 15:**
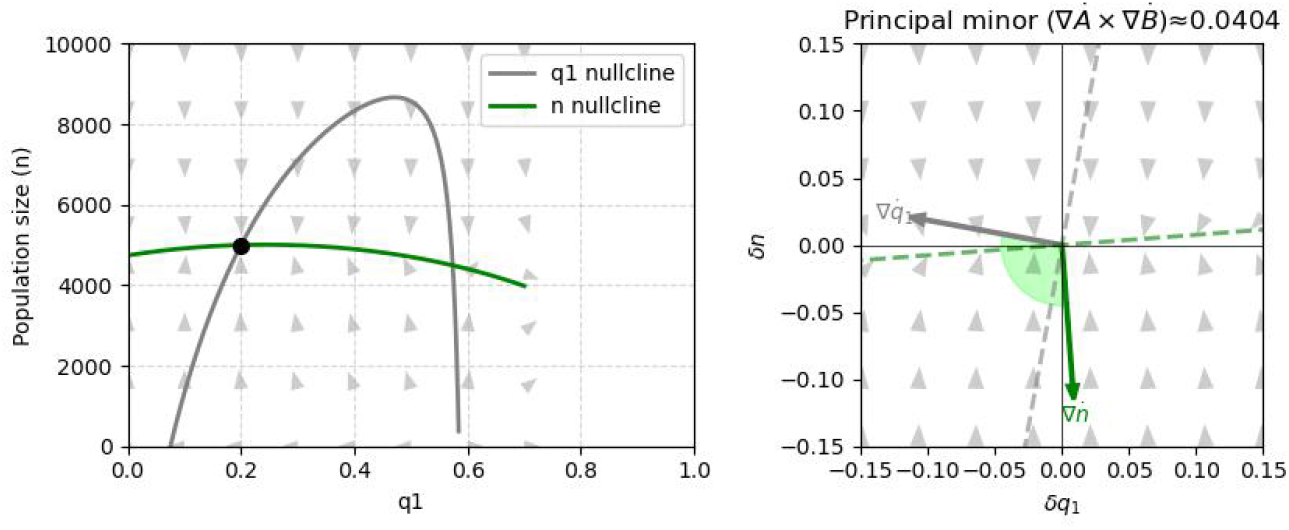
Example 1: Plot of the *q*_1_, *n* plane for the *q*_1_, *n* principal minor. The main intersection is stable in this plane. A secondary intersection appears as a proxy for the basin of attraction along the density manifold and the nullcline maximum marks the switch between its stable and unstable branches.

**Fig. 16:**
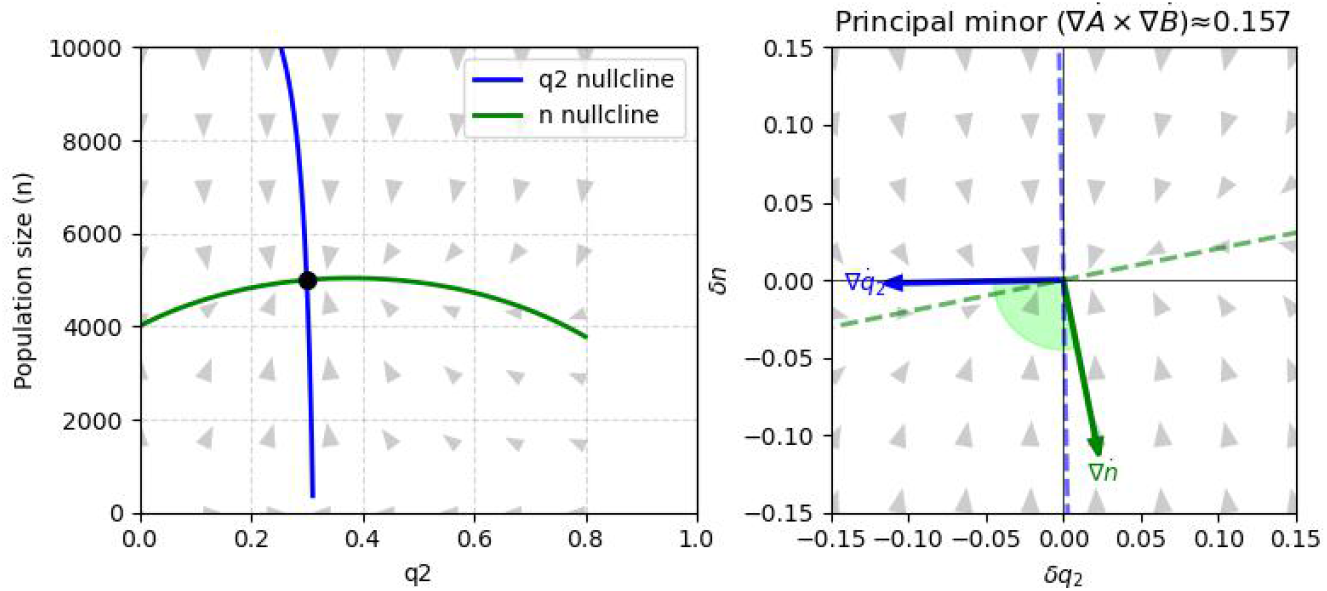
Example 1: Plot of the remaining “wall” plane corresponding to the *q*_2_, *n* principal minor. The flow and nullclines intersect at a point that is likewise stable when dynamics are restricted to this plane.

To further solidify the argument of stability, we examine the Jacobian at the fixed point:

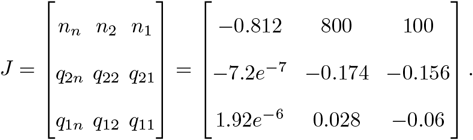

Its eigenvalues have negative real parts,

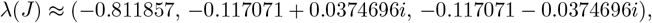

Stability can now be formally verified using the R-H conditions. At the fixed point, we compute

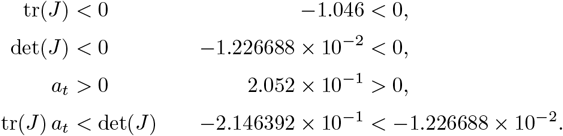

Since all Routh–Hurwitz conditions are satisfied, the fixed point is a *locally asymptotically stable sink*.

### 2. Stable vertex and boundary solutions; unstable interior solution

The trajectories that start from the interior of the simplex, where all three subpopulations exist initially, converge to stable boundary and corner solutions as can be seen in cell 30 from Appendix F. These solutions represent equilibrium states where either one or two of the strategies are eliminated, respectively. On the other hand, the interior solution is unstable and is a repeller of flow. The trajectories tend to move toward one of the stable boundary solutions, as the system evolves and eliminates one or more of the strategies. This dynamic highlights the dominance of boundary solutions over the interior in the long term.

The payoff matrices are :

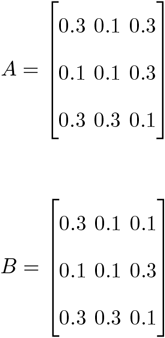

Rest points, trajectories, phase portraits and a heatmap are shown in Figures 17, 18, 19, 20 and 21. In figures 22 and 23 the structure of the nullclines is presented. Now we can focus on the geometric presentation of the stability conditions. Figures 24 and 25 show instability, while Figure 26 shows that the system is stable on this plane. Note that the overall sum of the principal minors is positive. However, the orientation of the Nash line inferred from the “floor” projection with the negative principal minor is co-directional with the density gradient. In effect, the three-dimensional condition fails, caused by a negative cross product in the *q*_1_, *q*_2_ plane.

**Fig. 17:**
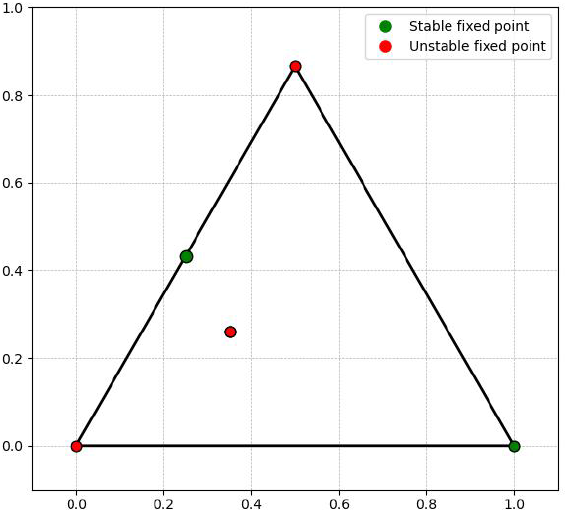
Example 2: Complete set of fixed points of the system, including both stable and unstable equilibria.

**Fig. 18:**
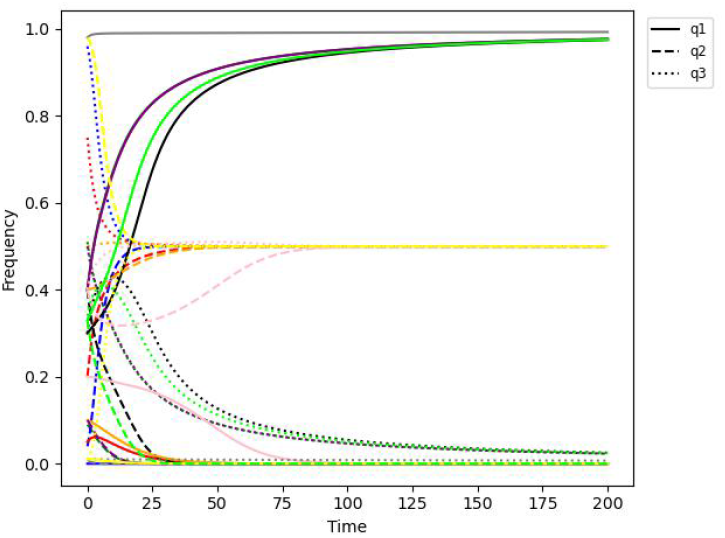
Example 2: Evolution of strategy frequencies over time.

**Fig. 19:**
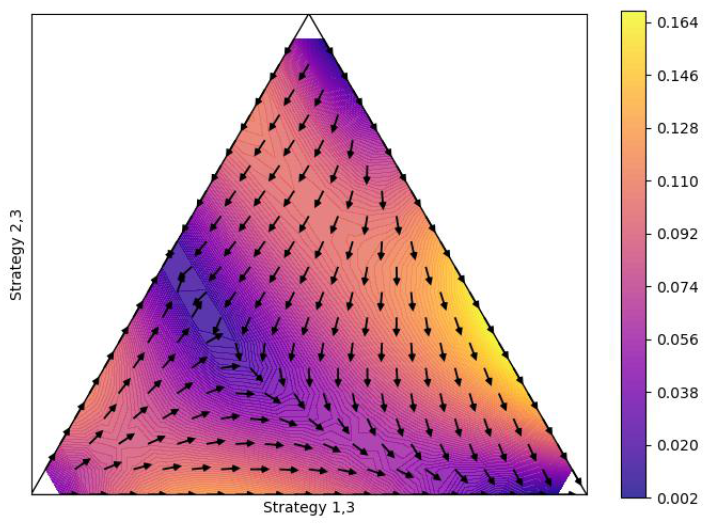
Example 2: Heatmap of the phase portrait illustrating the direction and speed of convergence.

**Fig. 20:**
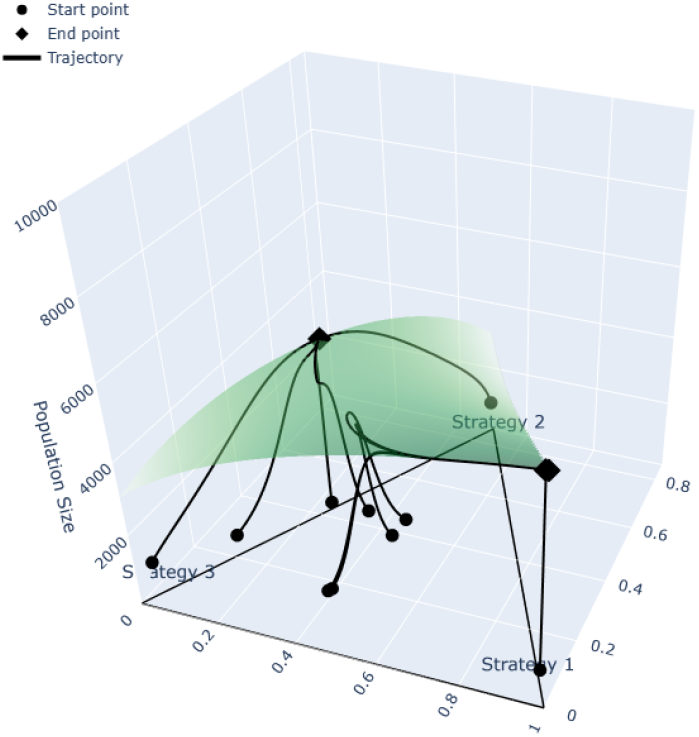
Example 2: Three-dimensional phase portrait with example trajectories, illustrating that trajectories originating from different regions are attracted towards boundary rest points.

**Fig. 21:**
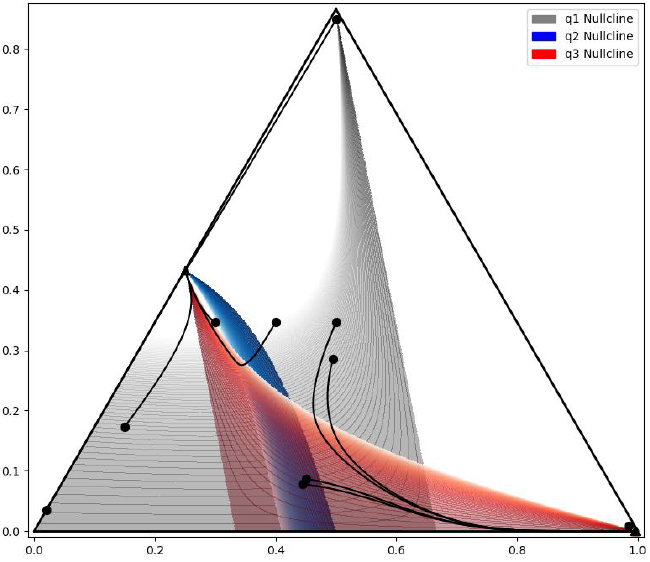
Example 2: Overhead view of trajectories superimposed on nullclines. The stable manifold of the interior saddle, which acts as a separatrix between boundary stable points, lies close to the *q*_1_ nullcline. The unstable manifold that attracts trajectories prior to convergence toward the stable point closely follows the shape of the *q*_3_ nullcline.

**Fig. 22:**
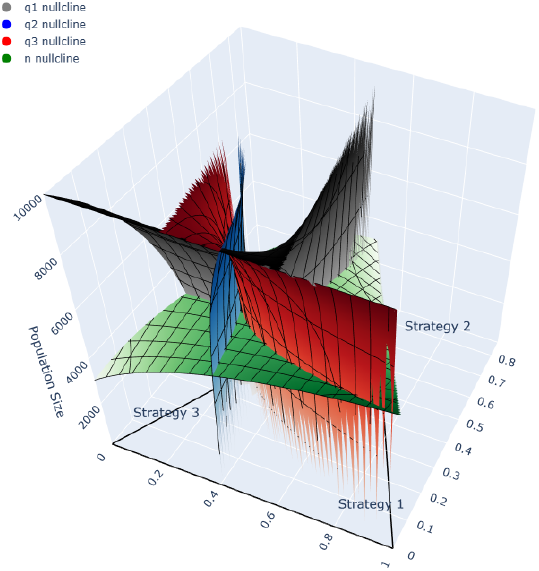
Example 2: 3D nullcline intersection.

**Fig. 23:**
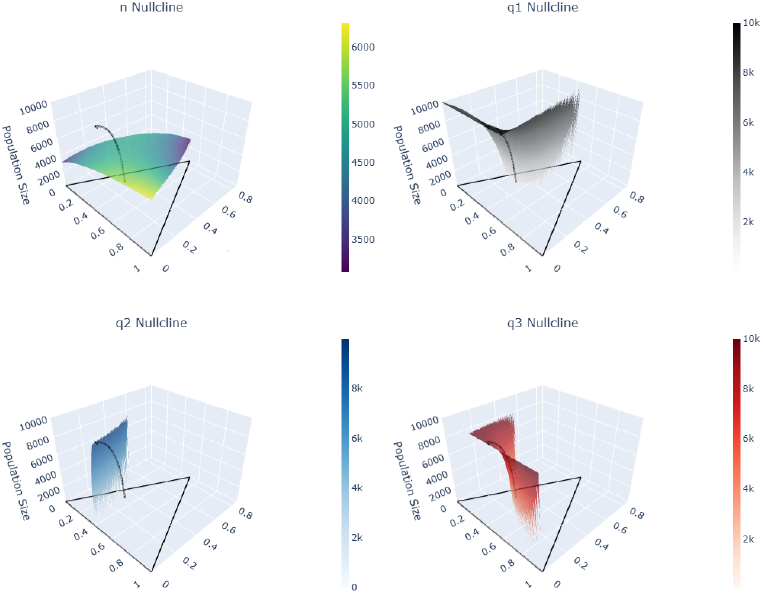
Example 2: 3D illustration of individual nullclines, with line of Nash Equilibria(from top left, clockwise): Population nullcline, *q*_1_ nullcline, *q*_3_ nullcline, *q*_2_ nullcline.

**Fig. 24:**
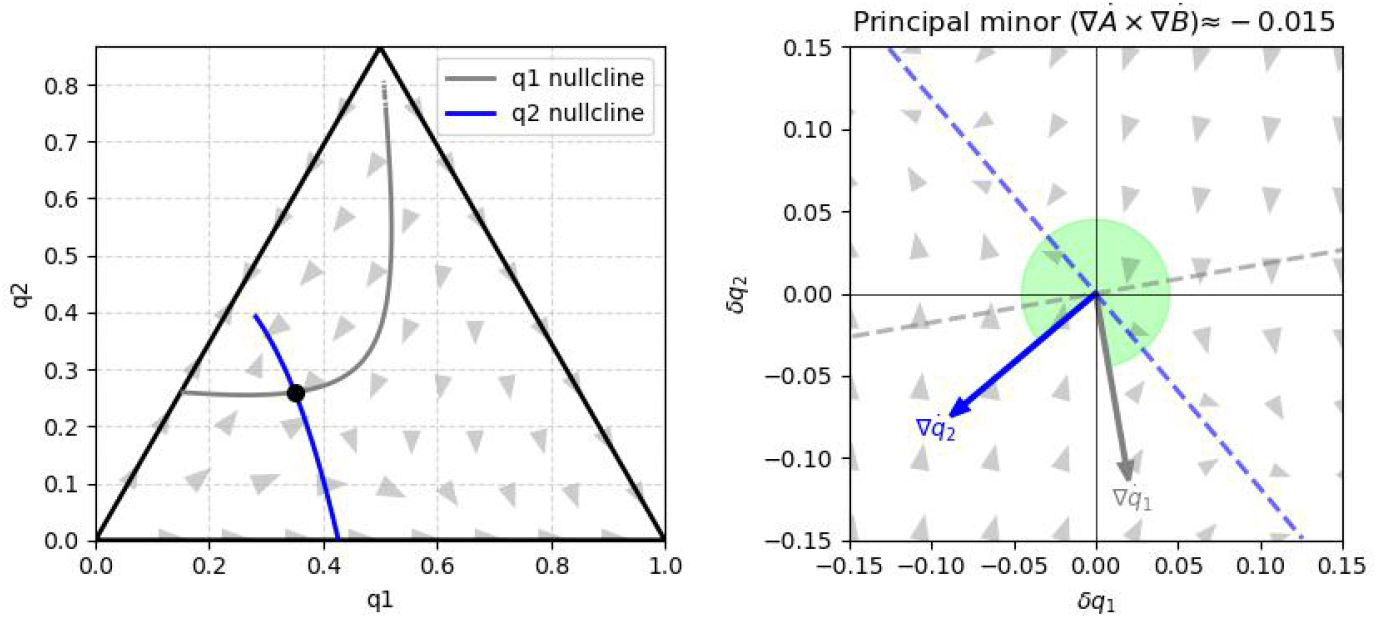
Example 2: Plot of the “floor” plane given by the *q*_1_, *q*_2_ principal minor. The intersection is unstable with saddle-type dynamics in this plane, implying a negative orientation of the Nash line. Consequently, the three-dimensional condition fails because the tangent to the Nash line is co-directional with the density gradient, although the sum of principal minors remains positive.

**Fig. 25:**
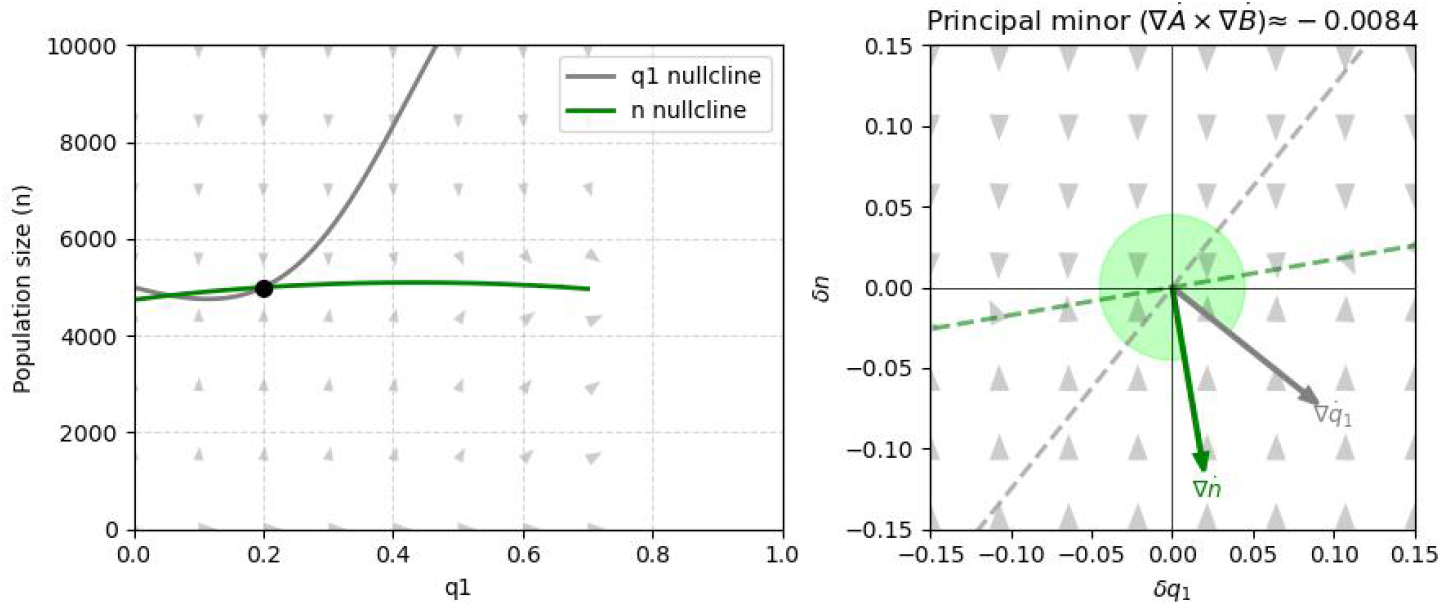
Example 2: Plot of the *q*_1_, *n* plane corresponding to the *q*_1_, *n* principal minor. The intersection is unstable in this plane, consistent with the dynamics observed in the “floor” plot, where the flow along the *q*_1_ axis is also unstable.

**Fig. 26:**
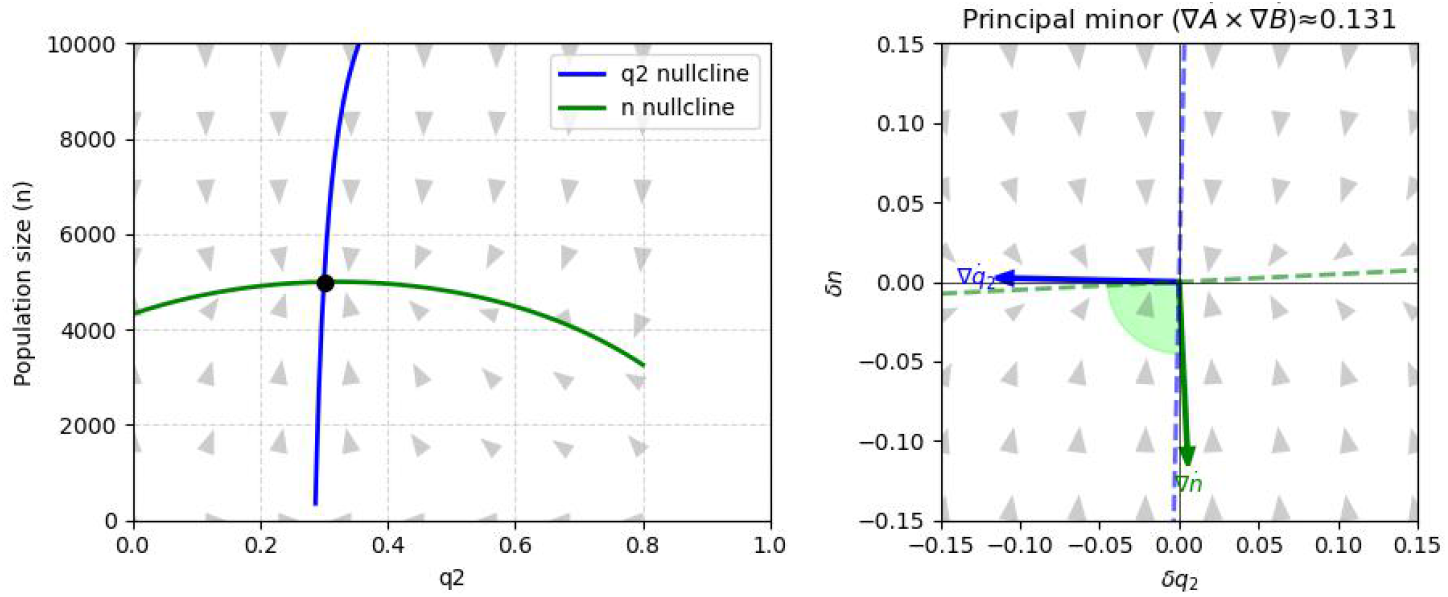
Example 2: Plot of the second “wall” plane defined by the *q*_2_, *n* cross product. In this plane, the intersection is stable, which is consistent with the flow in the “floor” plane showing the saddle dynamics.

At the interior fixed point we have, *q*_1_ = 0.2, *q*_2_ = 0.3, *q*_3_ = 0.5, *n* = 5000.

To further establish stability, we examine the Jacobian at the fixed point:

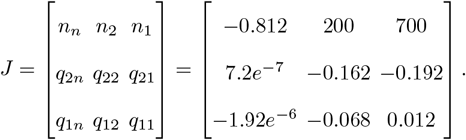

Stability can now be formally verified using the R-H conditions. At the fixed point, we compute

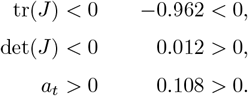

As the second condition is not satisfied, the interior fixed point is unstable.

### 3. Boundary solutions with stable and unstable fixed points

The trajectories which start from the interior of the simplex i.e. the ones which have all three subpopulations, converge to a boundary solution, thus eliminating one of the strategies. However, the trajectory which starts from the boundary i.e. which has only two subpopulations in the beginning, continues to stay on the same boundary solution and not escape to the strongly stable steady state. This is observed in cell 58 in Appendix F.

The payoff matrices are :

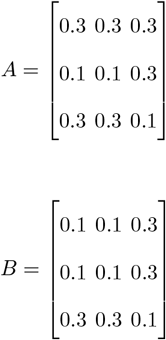

Consequently, we obtain the reduced relations as per (13) and (14) as follows

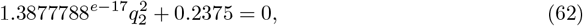

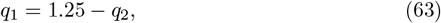

which do not reconcile with the general conditions established in Table 2. Hence, it confirms the absence of an interior fixed point. Figures 27, 28, 29, 30 and 31 show the rest points, trajectories and phase portraits. The structure of the nullclines is presented in Figures 32 and 33. The geometric interpretation of the stability conditions for the unstable boundary rest point is seen in Figures 34, 35 and 36. Respectively for the stable point, those conditions are verified in Figures 37, 38 and 39.

**Fig. 27:**
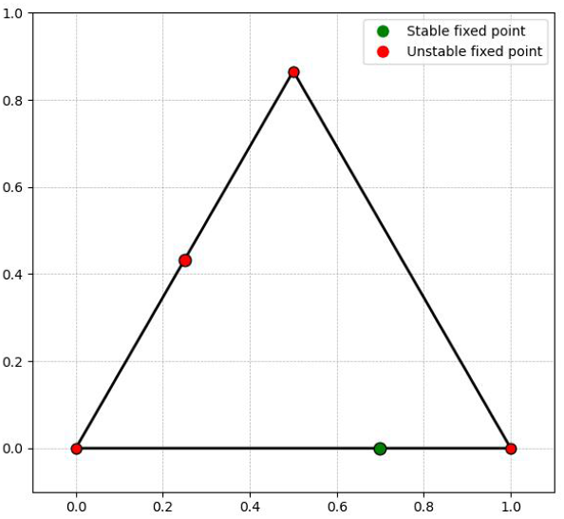
Example 3: Complete set of fixed points of the system, including both stable and unstable equilibria.

**Fig. 28:**
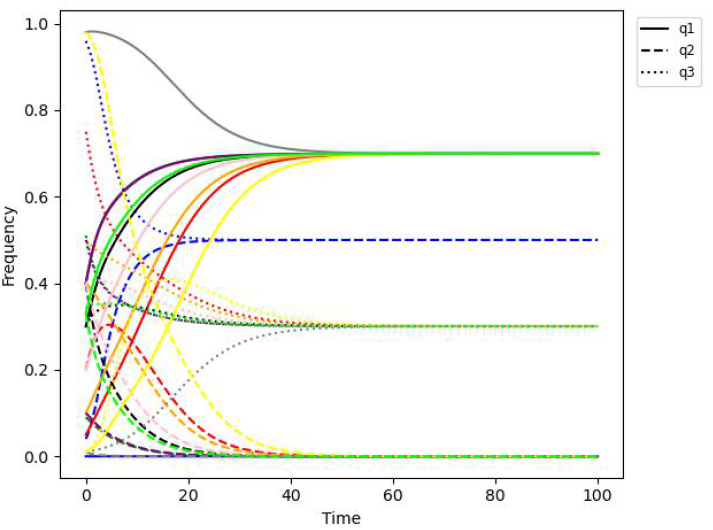
Example 3: Evolution of strategy frequencies over time.

**Fig. 29:**
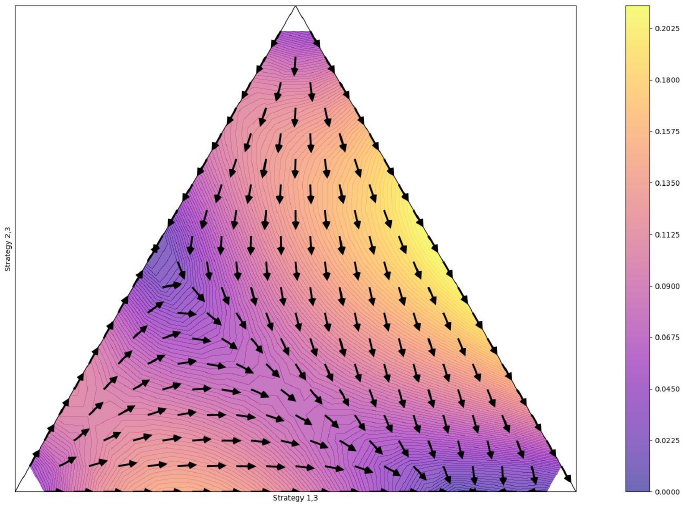
Example 3: Heatmap of the phase portrait illustrating the direction and speed of convergence.

**Fig. 30:**
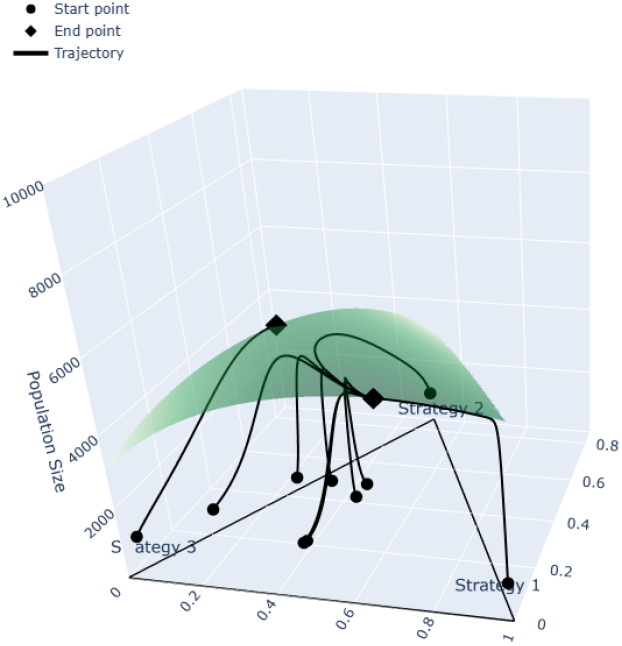
Example 3: 3-dimensional phase portrait with example trajectories, showing attraction toward the density nullcline and near it, motion along the heteroclinic orbit connecting the two rest points, followed by convergence to the stable rest point.

**Fig. 31:**
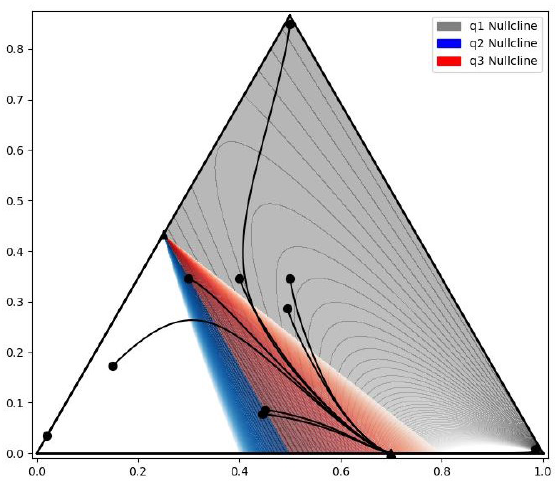
Example 3: Overhead view of trajectories superimposed on nullclines.

**Fig. 32:**
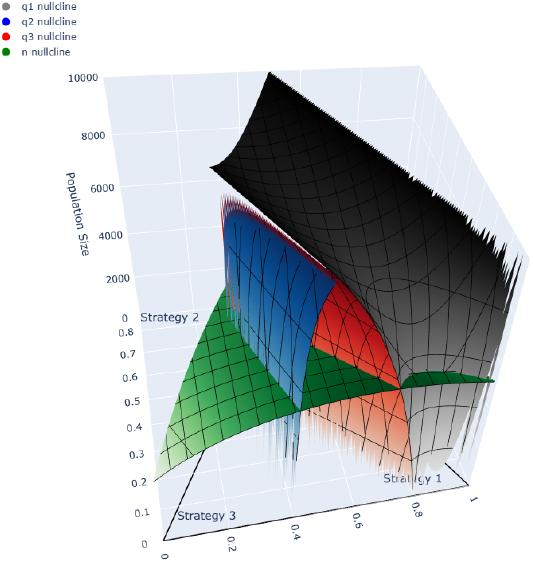
Example 3: 3D nullcline intersection.

**Fig. 33:**
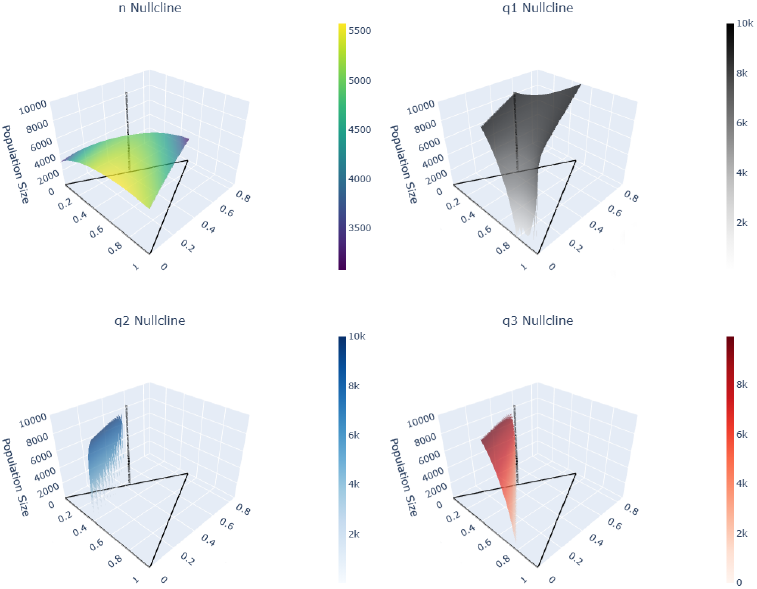
Example 3: 3D illustration of individual nullclines, with line of Nash Equilibria(from top left, clockwise): Population nullcline, *q*_1_ nullcline, *q*_3_ nullcline, *q*_2_ nullcline.

**Fig. 34:**
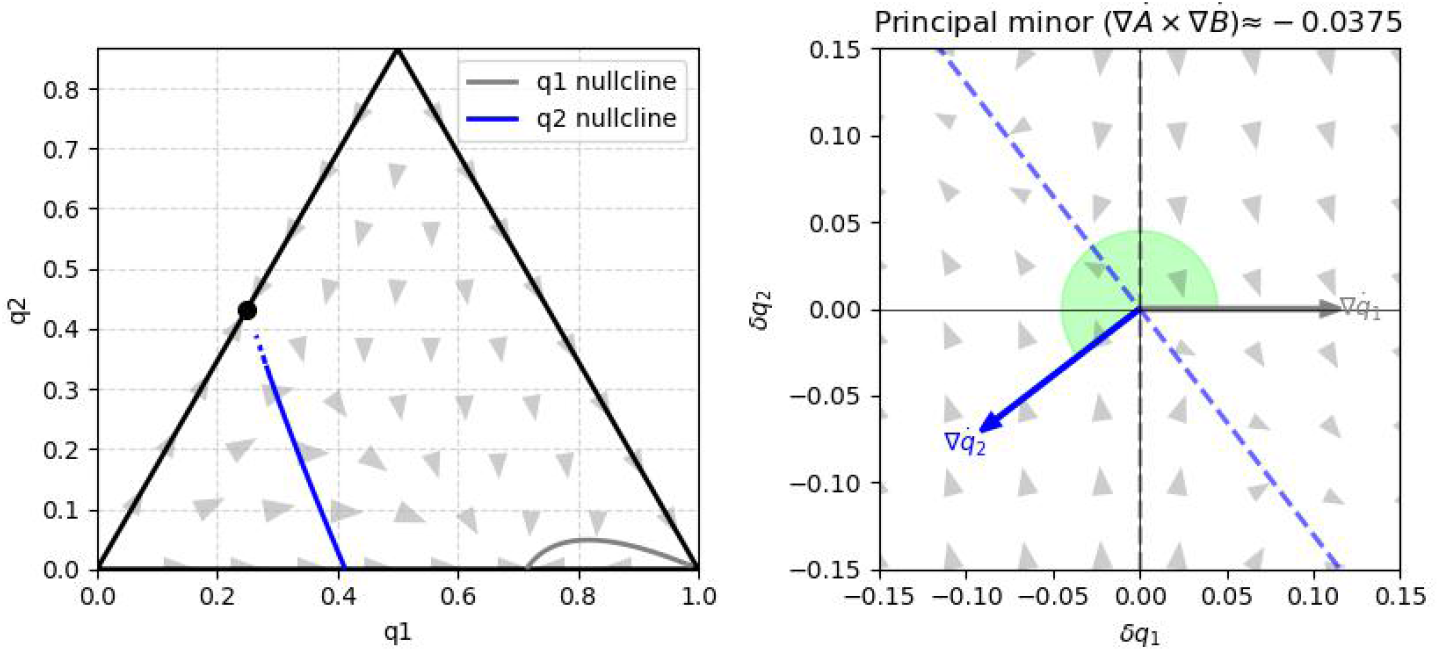
Example 3 (unstable rest point): The “floor” projection and the intersection of the internal *q*_2_ nullcline with trivial nullcline *q*_1_ = 0. The intersection is unstable.

**Fig. 35:**
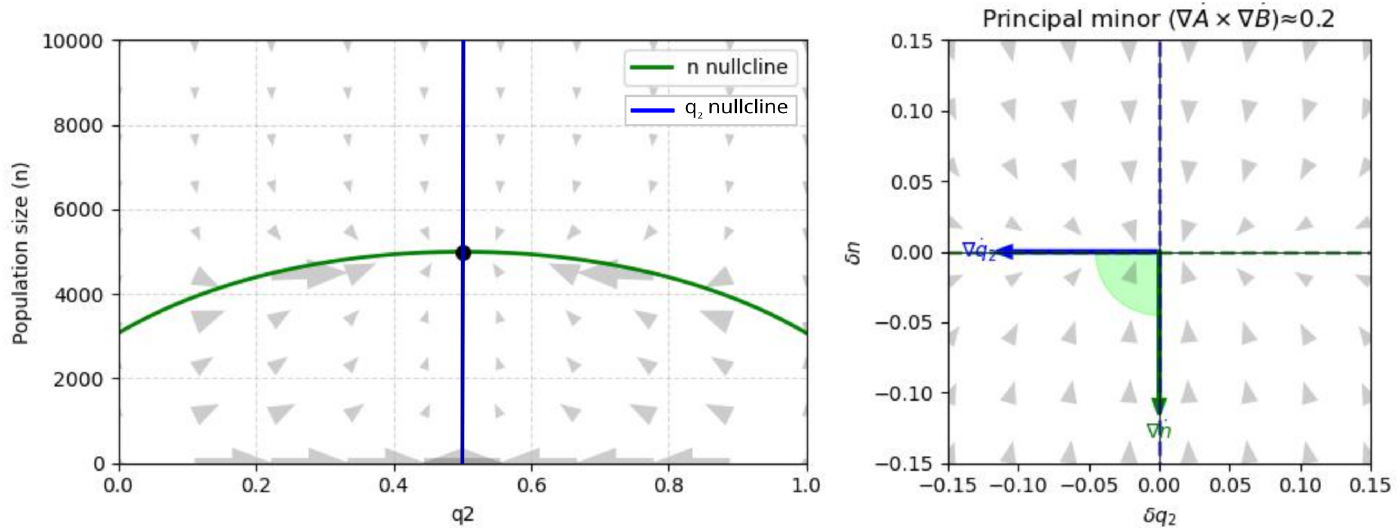
Example 3 (unstable rest point): The plots show the dynamics on the *q*_2_, *n* “wall”. We can see that in the absence of the strategy 1 (*q*_1_ = 0) the intersection is stable. However, the appearance of the rare mutant carrying the strategy 1 will shift the population towards the other rest point along the heteroclinic connection between them. The *q*_2_ nullcline is vertical since the fertility bracket is zero at the boundary, which means that there are no fertility differences between strategy 1 and 3. Note that the flow in that plane is symmetric.

**Fig. 36:**
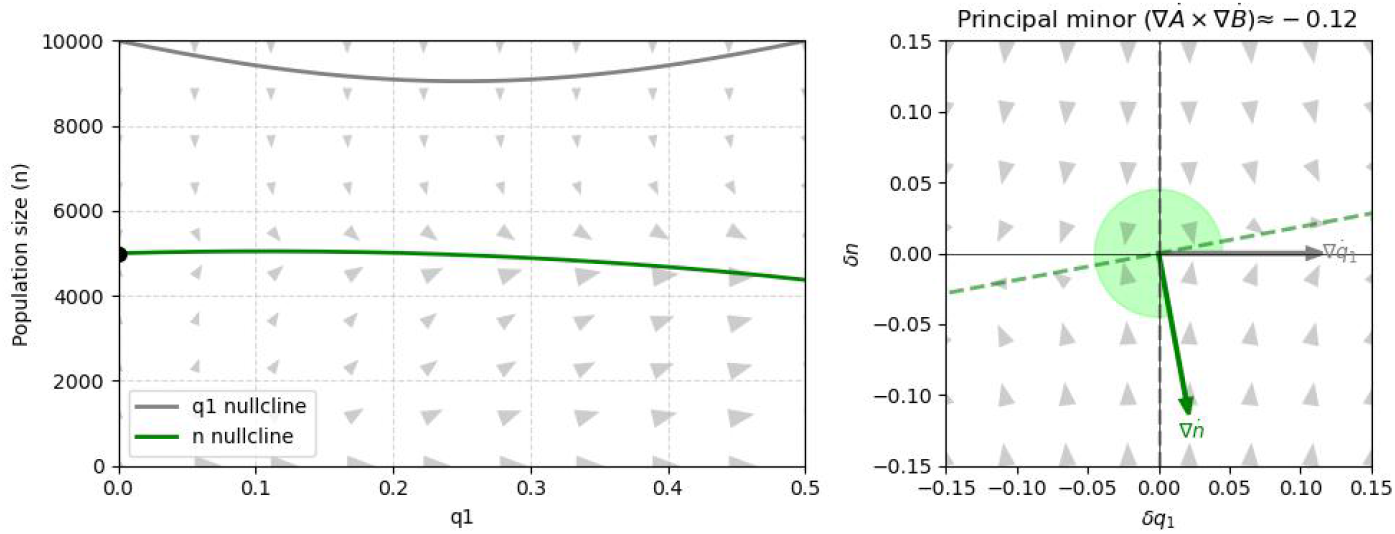
Example 3 (unstable rest point): Plot of the *q*_1_, *n* plane showing the intersection of the trivial nullcline *q*_1_ = 0 with the density nullcline. The intersection is unstable, illustrating escape dynamics driven by invasion of the small amount of strategy-1 individuals.

**Fig. 37:**
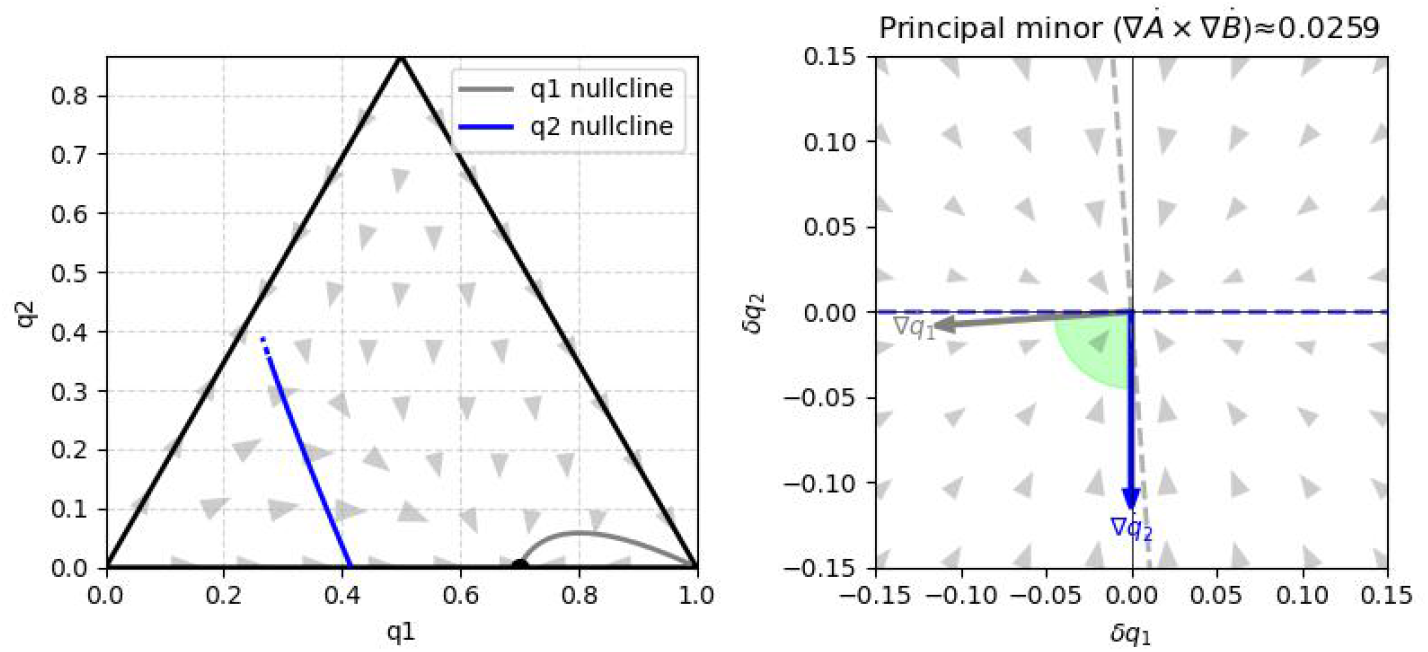
Example 3 (stable rest point): Plot shows the “floor” intersection of the trivial nullcline *q*_2_ = 0 with nontrivial *q*_1_ nullcline (nontrivial *q*_2_ nullcline is also shown). The intersection is stable.

**Fig. 38:**
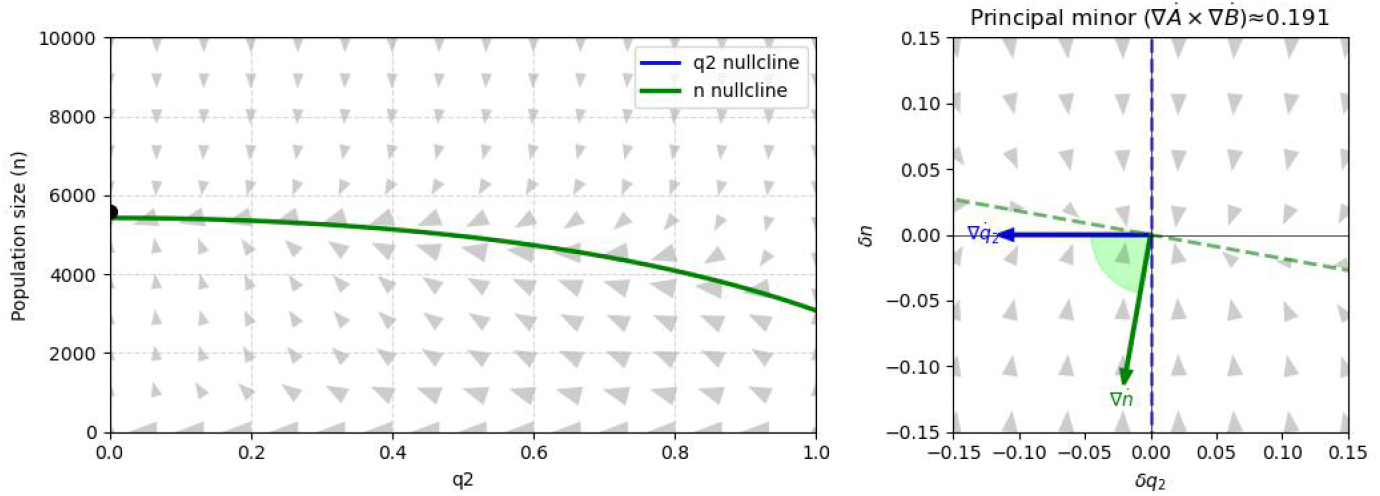
Example 3 (stable rest point): The plot of the second “wall” plane described by *q*_2_, *n* cross product. In this plane, the intersection of trivial *q*_2_ nullcline and density nullcline is stable, which is consistent with the flow in the “floor” plane showing the attraction.

**Fig. 39:**
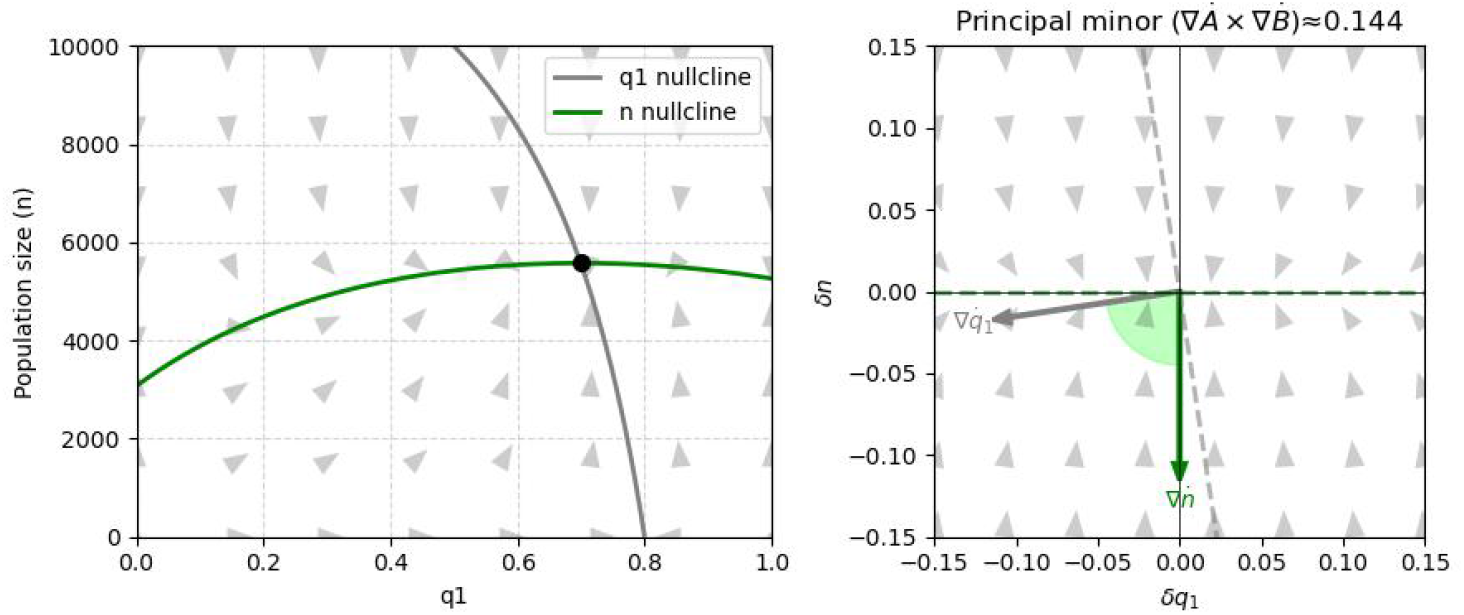
Example 3 (stable rest point): The plot of the *q*_1_, *n* plane described by *q*_1_, *n* principal minor. The intersection is stable in this plane. We can see that in the case of boundary rest points, the respective “wall” condition is equivalent to the three dimensional condition (see Figure 32).

## 6 Discussion

General matrix games with eco-evolutionary feedback are usually studied using only two strategies, for example cooperators and defectors [34] or Hawks and Doves [7, 24, 26]. In this work, the demographic game proposed in [24] is generalized to three strategies and enhanced with new methods for verifying stability and simplifying calculations. Expanding the strategy space in this way reveals richer and more biologically realistic dynamics that cannot be captured in two-strategy systems.

We have applied the mechanistic framework to a subset of closed-form solutions constrained by the exogenous parameter *ω*, which ensures analytical tractability. Through this extension, we are able to visualize the symbolic structure of how the dynamics unfold, while examining the interaction of strategies to attain steady-state compositions under a fixed population size. The key results are summarized as necessary conditions that must hold for the existence of an interior fixed point. Also, the extensive table of dynamics achieved by the constraint helps in narrowing the payoff matrix structure, for which the trajectories verifiably move between the fixed points in the phase space.

A central component of our contribution is a geometric perspective on stability analysis. Each element of the Routh-Hurwitz conditions is interpreted through the intersections and relative orientation of nullclines in the phase space, allowing regions of stability and instability to be identified directly from their geometric configuration. This provides an intuitive connection between the stability of a rest point and the characteristic flow patterns in its neighbourhood. The nullcline-based approach can also substantially reduce computational effort. In many cases, stability can be inferred solely from the ordering of nullclines in two-dimensional projections and from the one-dimensional intersection (a Nash line in our replicator system) formed by two nullclines crossing the third, stationary-density nullcline. Thus, stability properties can often be read directly from the shapes and positions of nullclines in the figures. When the system is more intricate, the analysis typically reduces to relatively simple algebraic inequalities.

In addition, the geometric method allows for deeper insight into the underlying mechanics. It highlights the specific mechanisms responsible for stability loss, such as changes in the ordering of stable nullclines or the loss of stability of individual nullclines. These features are reflected in the angles and orientations of the nullcline surfaces along the coordinate axes. Our approach underscores the fundamental role of nullclines in the analysis of rest-point stability and other qualitative properties of dynamical systems. Nullclines generally delineate regions of growth and decline and constitute surfaces of trajectory extrema (minima or maxima). Their relevance extends beyond stability analysis as nullcline intersections can also organize more complex behaviour, including chaotic dynamics that circulate around them [33]. Within this framework, we make use of cross products and their geometric interpretation to connect with established stability conditions for two-dimensional models [26], while substantially extending earlier nullcline and gradient-based methodologies developed for planar systems [35]. In particular, we introduce the use of gradient projections onto respective complementary nullclines enabling stability inference directly from the flow along nullclines, together with cross-product geometry capturing stability information from the ordering of nullclines. Overall, our framework provides a more rigorous and comprehensive methodology tailored to three-dimensional models. While the geometric method clarifies whether a rest point is stable, one may still ask why it is stable. Addressing this question requires quantitative relationships among model parameters. Here, vector calculus proves useful: it eliminates redundancies and focuses attention on the focal payoffs of each strategy, revealing the algebraic structure of the underlying mechanistic laws.

Our numerical examples illustrate the interplay between selection mechanisms and ecological factors, including the concavity or convexity of the stationary density surface. They also emphasize the importance of heteroclinic orbits connecting rest points. In particular, heteroclinic connections between boundary and interior equilibria are typically slower and tend to attract nearby trajectories. Biologically, they can be interpreted as the transients from a monomorphic or two-strategy equilibrium to a three-strategy system equilibrium resulting from the invasion of a rare mutant.

## Acknowledgments

This work is generously supported by the studentship awarded to Manjyot Singh Bedi by City St. George’s, University of London, for which we are very grateful. Krzysztof Argasinski was supported by grant OPUS 2020/39/B/NZ8/03485 awarded by Polish National Science Center.

## Declarations

The authors declare no conflict of interest.

## Appendix A Reduction of dimension

In order to understand the interplay between the two independent strategies, we reduce the system to two dimensions using the following compression :

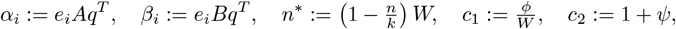

where *e*_*i*_ is the unit vector corresponding to strategy *i*. Introducing the shorthand,

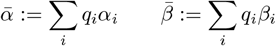

The equilibrium conditions are then

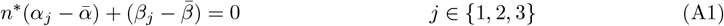

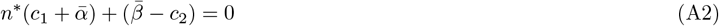

The value of the logistic suppression coefficient will be

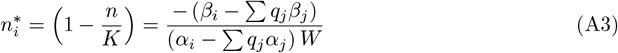

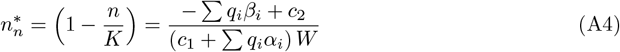

Then the nullclines will be:

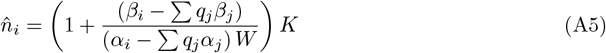

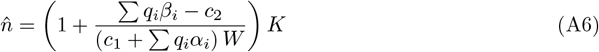

By adding (A1) for *j* = 1 to (A2), and subtracting the cases *j* = 2, 3 from the case *j* = 1, we obtain:

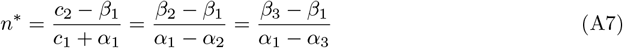

On eliminating *n*^∗^, the unified condition (A7) decomposes naturally into two parts:

### 1. Background balance

consistency between ecological parameters (*c*_1_, *c*_2_) and the strategy payoffs, given by

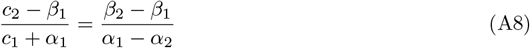

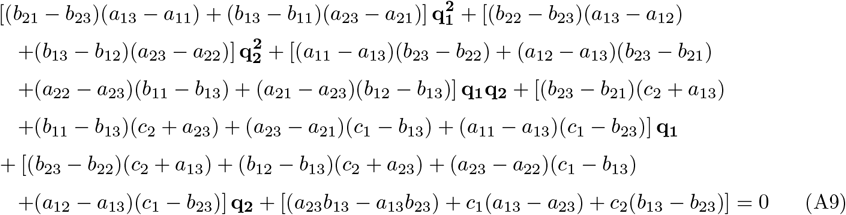

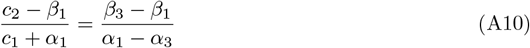

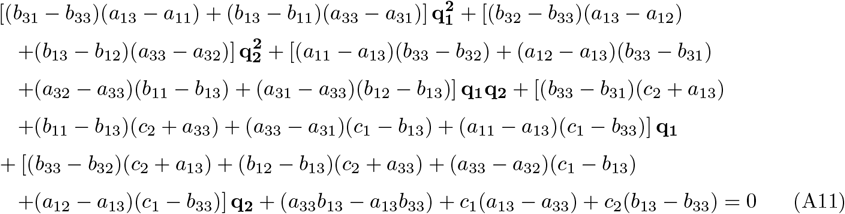

#### 2. Indifference of strategies

mutual equality of the focal payoffs, independent of the background constants,

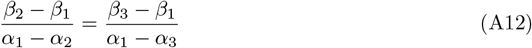

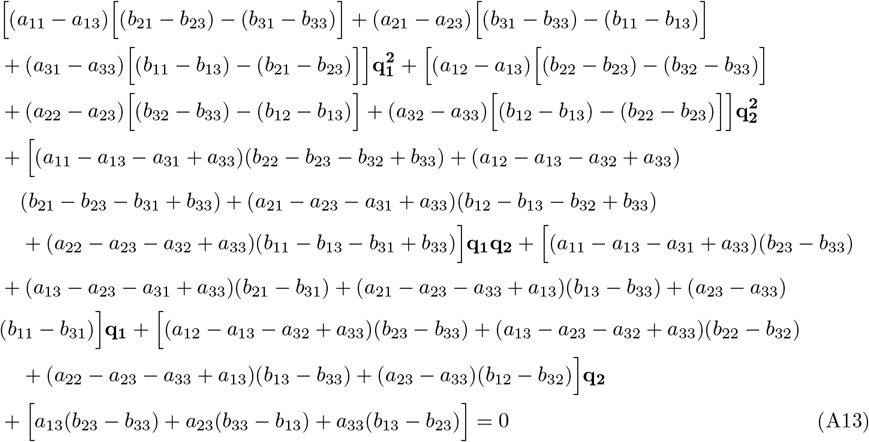

Using auxiliary notation, our working set of equations, (A9), (A11) and (A13), are represented as:

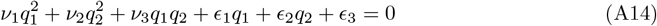

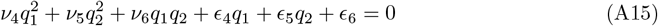

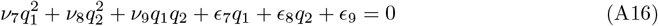

This notation can be referenced from Table A1.

## Appendix B Constrained Roots

**Table A1:** Definitions of coefficients *ν*_*i*_ and *ϵ*_*i*_.

|  |  |
| --- | --- |
| $\nu_1$ | $(b_{21} - b_{23})(a_{13} - a_{11}) + (b_{13} - b_{11})(a_{23} - a_{21})$ |
| $\nu_2$ | $(b_{32} - b_{33})(a_{13} - a_{12}) + (b_{13} - b_{12})(a_{23} - a_{22})$ |
| $\nu_3$ | $(a_{11} - a_{13})(b_{23} - b_{22}) + (a_{12} - a_{13})(b_{23} - b_{21}) + (a_{22} - a_{23})(b_{11} - b_{13}) + (a_{21} - a_{23})(b_{12} - b_{13})$ |
| $\nu_4$ | $(b_{31} - b_{33})(a_{13} - a_{11}) + (b_{13} - b_{11})(a_{33} - a_{31})$ |
| $\nu_5$ | $(b_{32} - b_{33})(a_{13} - a_{12}) + (b_{13} - b_{12})(a_{33} - a_{32})$ |
| $\nu_6$ | $(a_{11} - a_{13})(b_{33} - b_{32}) + (a_{12} - a_{13})(b_{33} - b_{31}) + (a_{32} - a_{33})(b_{11} - b_{13}) + (a_{31} - a_{33})(b_{12} - b_{13})$ |
| $\nu_7$ | $(a_{11} - a_{13})[(b_{21} - b_{23}) - (b_{31} - b_{33})] + (a_{21} - a_{23})[(b_{31} - b_{33}) - (b_{11} - b_{13})] + (a_{31} - a_{33})[(b_{11} - b_{13}) - (b_{21} - b_{23})]$ |
| $\nu_8$ | $(a_{12} - a_{13})[(b_{22} - b_{23}) - (b_{32} - b_{33})] + (a_{22} - a_{23})[(b_{32} - b_{33}) - (b_{12} - b_{13})] + (a_{32} - a_{33})[(b_{12} - b_{13}) - (b_{22} - b_{23})]$ |
| $\nu_9$ | $(a_{11} - a_{13} - a_{31} + a_{33})(b_{22} - b_{23} - b_{32} + b_{33}) + (a_{12} - a_{13} - a_{32} + a_{33})(b_{21} - b_{23} - b_{31} + b_{33})$<br>$+ (a_{21} - a_{23} - a_{31} + a_{33})(b_{12} - b_{13} - b_{32} + b_{33}) + (a_{22} - a_{23} - a_{32} + a_{33})(b_{11} - b_{13} - b_{31} + b_{33})$ |
| $\epsilon_1$ | $(b_{33} - b_{31})(c_2 + a_{13}) + (b_{11} - b_{13})(c_2 + a_{23}) + (a_{23} - a_{21})(c_1 - b_{13}) + (a_{11} - a_{13})(c_1 - b_{23})$ |
| $\epsilon_2$ | $(b_{23} - b_{22})(c_2 + a_{13}) + (b_{12} - b_{13})(c_2 + a_{23}) + (a_{23} - a_{22})(c_1 - b_{13}) + (a_{12} - a_{13})(c_1 - b_{23})$ |
| $\epsilon_3$ | $(a_{23}b_{13} - a_{13}b_{23}) + c_1(a_{13} - a_{23}) + c_2(b_{13} - b_{23})$ |
| $\epsilon_4$ | $(b_{33} - b_{31})(c_2 + a_{13}) + (b_{11} - b_{13})(c_2 + a_{33}) + (a_{33} - a_{31})(c_1 - b_{13}) + (a_{11} - a_{13})(c_1 - b_{33})$ |
| $\epsilon_5$ | $(b_{33} - b_{32})(c_2 + a_{13}) + (b_{12} - b_{13})(c_2 + a_{33}) + (a_{33} - a_{32})(c_1 - b_{13}) + (a_{12} - a_{13})(c_1 - b_{33})$ |
| $\epsilon_6$ | $(a_{33}b_{13} - a_{13}b_{33}) + c_1(a_{13} - a_{33}) + c_2(b_{13} - b_{33})$ |
| $\epsilon_7$ | $(a_{11} - a_{13} - a_{31} + a_{33})(b_{23} - b_{33}) + (a_{13} - a_{23} - a_{31} + a_{33})(b_{21} - b_{31})$<br>$+ (a_{21} - a_{23} - a_{33} + a_{13})(b_{13} - b_{33}) + (a_{23} - a_{33})(b_{11} - b_{31})$ |
| $\epsilon_8$ | $(a_{12} - a_{13} - a_{32} + a_{33})(b_{23} - b_{33}) + (a_{13} - a_{23} - a_{32} + a_{33})(b_{22} - b_{32})$<br>$+ (a_{22} - a_{23} - a_{33} + a_{13})(b_{13} - b_{33}) + (a_{23} - a_{33})(b_{12} - b_{32})$ |
| $\epsilon_9$ | $a_{13}(b_{23} - b_{33}) + a_{23}(b_{33} - b_{13}) + a_{33}(b_{13} - b_{23})$ |

Let

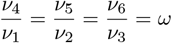

So,

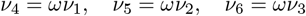

Substituting these in (A11) leads to,

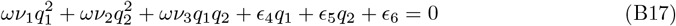

Subtracting *ω*(A9) from A11

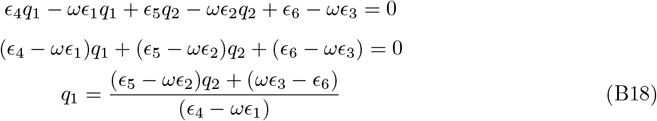

Substituting in (A9)

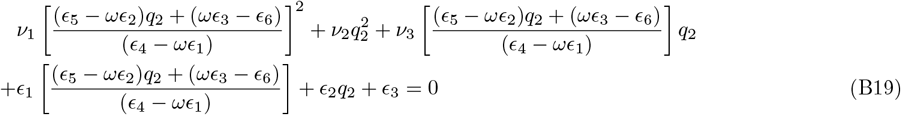

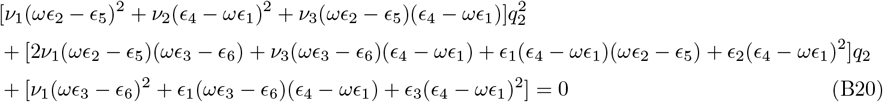

## Appendix C Modified final R-H Condition

R-H criterion is completed by,

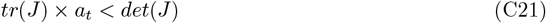

Explicitly, in our system’s notation, this is written as

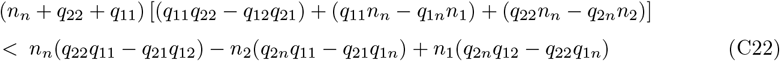

On expansion, we get

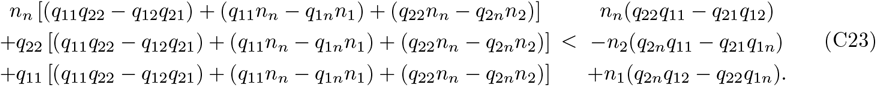

Canceling common terms on either side of the inequality,

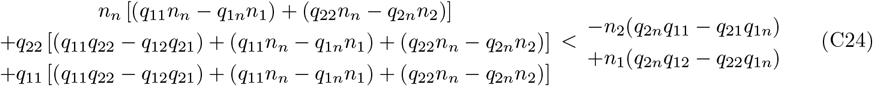

On further simplification, we get

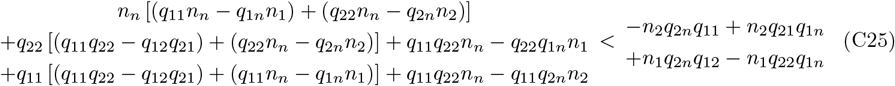

This leads to further elimination as

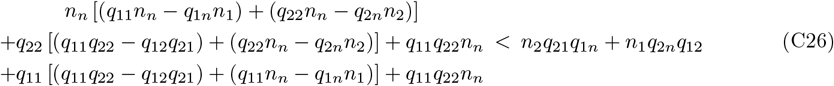

Hence, our final expression is

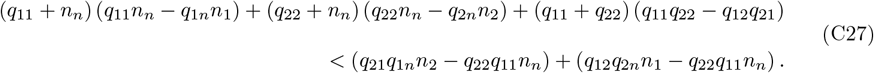

## Appendix D Intersections and Slopes of Nullclines

From Figure 6 we have that *ϕ* denotes the angle between any two gradients and when they are projected on one of the planes, the superscript of the orthogonal axis is added. The remaining angles on the projected plane, are denoted by *α*_*i*_, where *i* represents the focal axis which completes the angle. In addition, *β* is the angle between 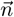 gradient and the vector tangent to the Nash line resulting from the cross product of 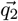 and 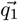.

The slope of gradient describing the variable *A* with respect to its own axis is

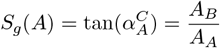

But for the nullcline slope along the focal axis we need zero directional derivative,

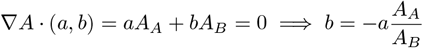

Thus the slope of the nullcline of the equation describing the variable *A* (focal variable is described by subscript) with respect to the axis described by superscript is

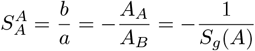

which is the negative reciprocal of the slope of the gradient.

Also, the slope of the nullcline projection along the opposite axis is given as,

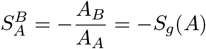

Since 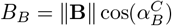 and 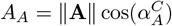, we have:

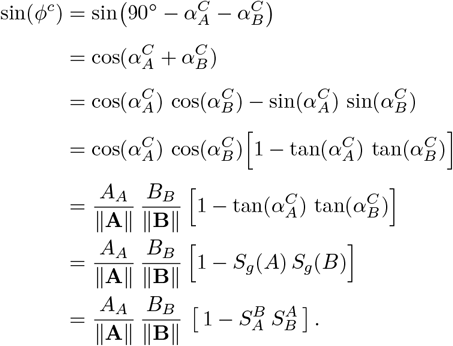

Thus the Two-dimensional condition can be written as,

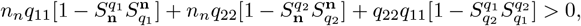

along with the cross-dimensional condition,

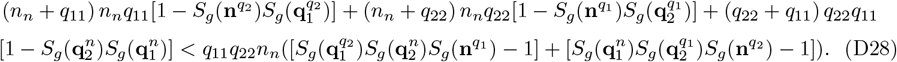

which for *q*_11_*q*_22_*n*_*n*_ < 0 (*q*_11_*q*_22_*n*_*n*_ > 0) can be simplified to the form:

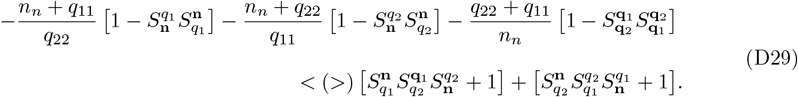

## Appendix E Mechanics of Gradient Directional Decomposition

### E.1 Three-dimensional

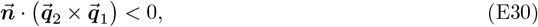

where

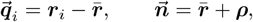

and 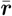 denotes the weighted average of the three focal payoff *gradient* vectors ***r***_1_, ***r***_2_, ***r***_3_:

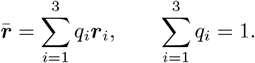

Expanding the cross product,

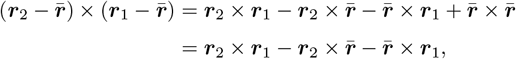

since 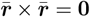. Therefore,

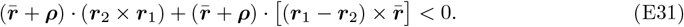

Now, note that

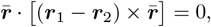

because it is a scalar triple product with two identical vectors. Hence, only the ***ρ*** part contributes, giving

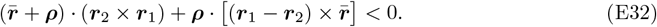

Equivalently,

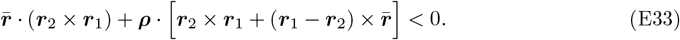

Since 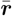 is a weighted average of ***r***_1_, ***r***_2_, ***r***_3_, we can write

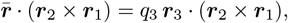

because ***r***_1_ · (***r***_2_ × ***r***_1_) = ***r***_2_ · (***r***_2_ × ***r***_1_) = 0.

For the second term, observe that

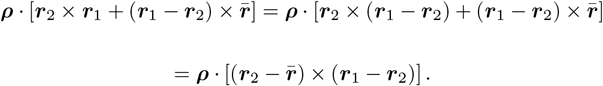

Now,

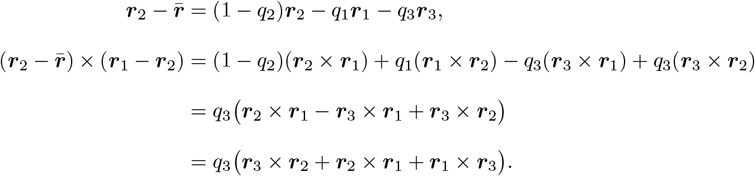

Therefore, the inequality can finally be expressed as

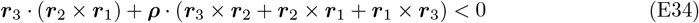

#### E.2 Two-dimensional

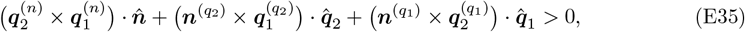

where, for each projection index *j* ∈ {*n, q*_1_, *q*_2_},

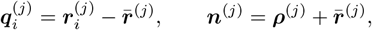

and 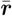 denotes the weighted average of the three focal payoff gradients:

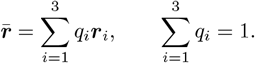

In the following intermediate steps we work in a fixed projection plane and suppress the superscripts ^(*j*)^ for readability. Expanding the first term,

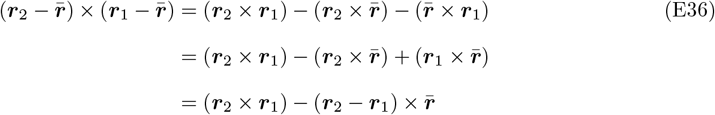

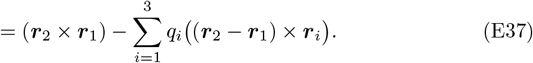

Using (***r***_2_ − ***r***_1_) × ***r***_1_ = ***r***_2_ × ***r***_1_ and (***r***_2_ − ***r***_1_) × ***r***_2_ = ***r***_2_ × ***r***_1_, we obtain

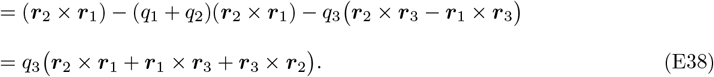

Restoring the (*n*)-superscripts, we obtain

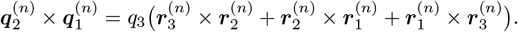

Similarly, the second term expands to

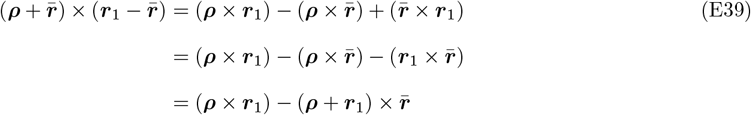

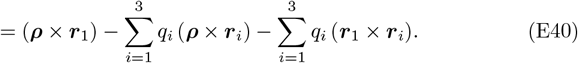

Since ***r***_1_ × ***r***_1_ = **0**,

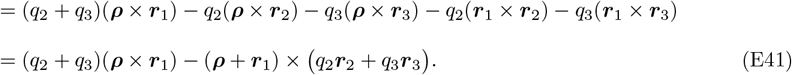

Restoring the (*q*_2_)-superscripts,

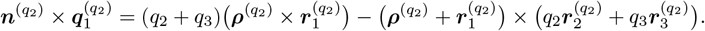

Similarly, the third term gives

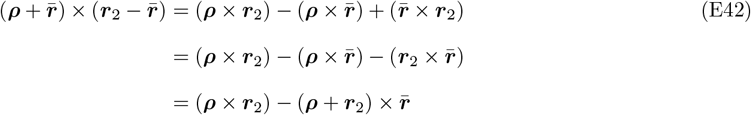

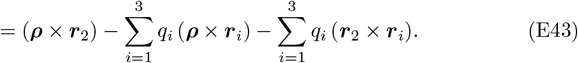

With ***r***_2_ × ***r***_2_ = **0**,

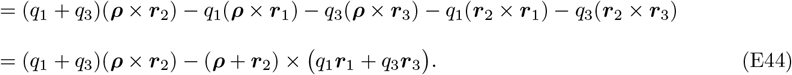

Restoring the (*q*_1_)-superscripts,

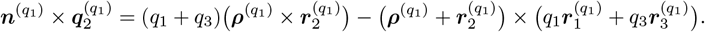

Therefore, the inequality can finally be expressed as

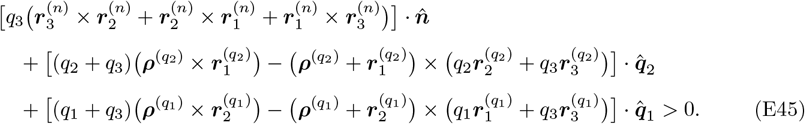

#### E.3 One-dimensional

We consider the replicator equation for strategy 1:

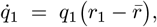

where the scalar average payoff is

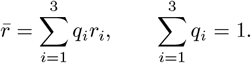

We compute the partial derivative 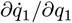, denoted *q*_11_:

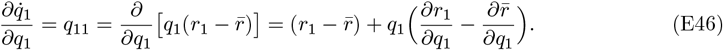

The partial derivative of the average payoff with respect to *q*_1_ is

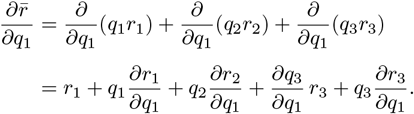

Since *q*_3_ = 1 − *q*_1_ − *q*_2_ we have ∂*q*_3_/∂*q*_1_ = −1. Hence

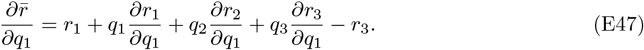

Using (E46) and (E47), we obtain

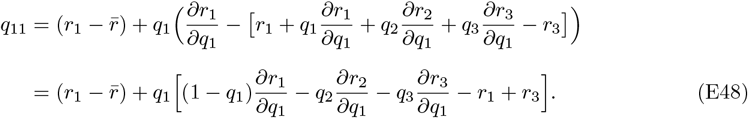

In our model the focal payoff of strategy *i* is composed of a fertility and a survival component:

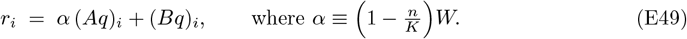

As the payoffs are linear in *q*,

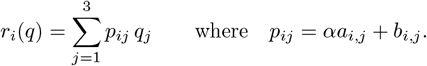

Using *q*_3_ = 1 − *q*_1_ − *q*_2_, we can write

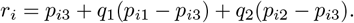

Substituting these constants into (E48) above gives

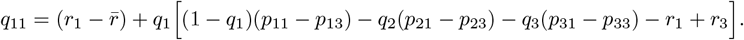

Evaluating at an interior equilibrium where 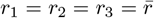, both 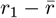 and −*r*_1_ + *r*_3_ vanish and we get

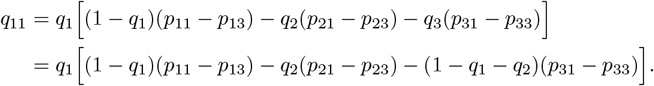

So we have

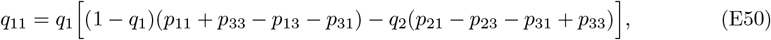

which becomes

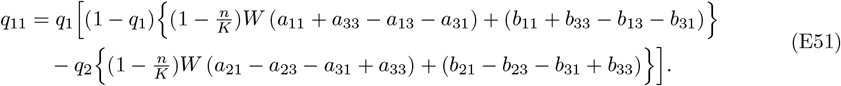

Similarly, for the second strategy we have

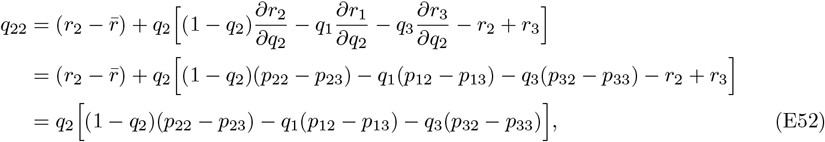

and hence

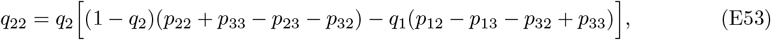

which becomes

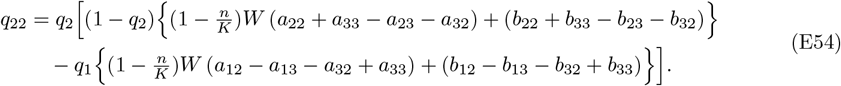

Since *r*_*i*_ is linear in *q*, the partial derivative with respect to *q*_*j*_ is

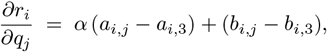

where *a* and *b* are entries of the fertility and survival matrices *A* and *B*.

For the density equation we have

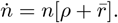

Then

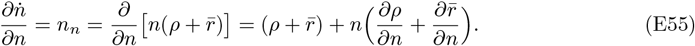

The partial derivative of the average payoff with respect to *n* is

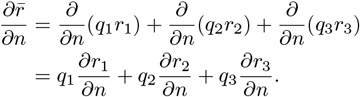

From (E49), we have

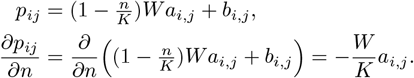

Using

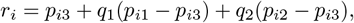

we obtain

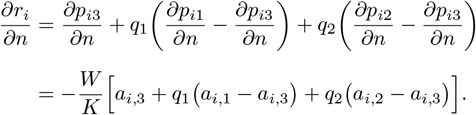

Thus

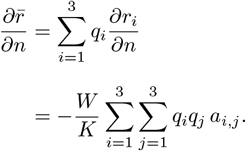

Also,

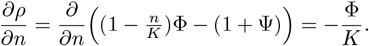

Evaluating at the non-trivial equilibrium, we have 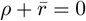, so

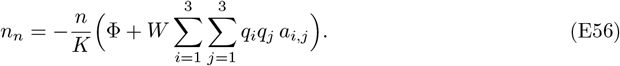

Therefore, the trace condition tr(*J*) < 0 can be written as

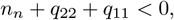

or explicitly,

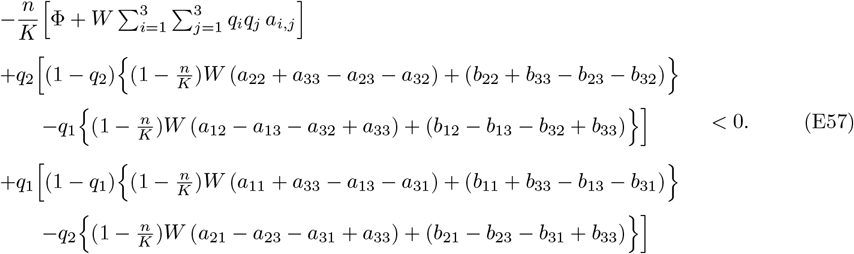

## Appendix F Table of Dynamics - Nullclines and Trajectories

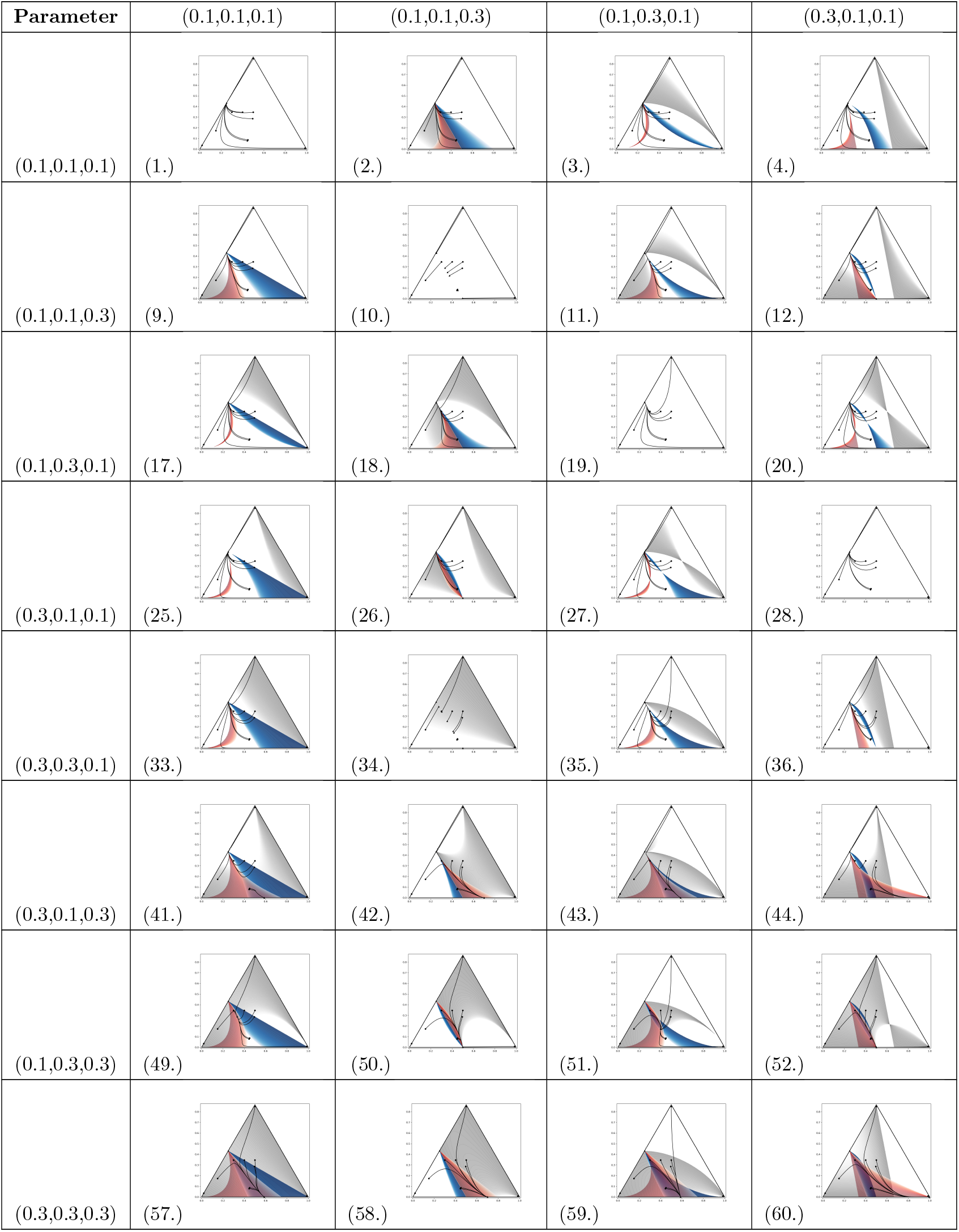
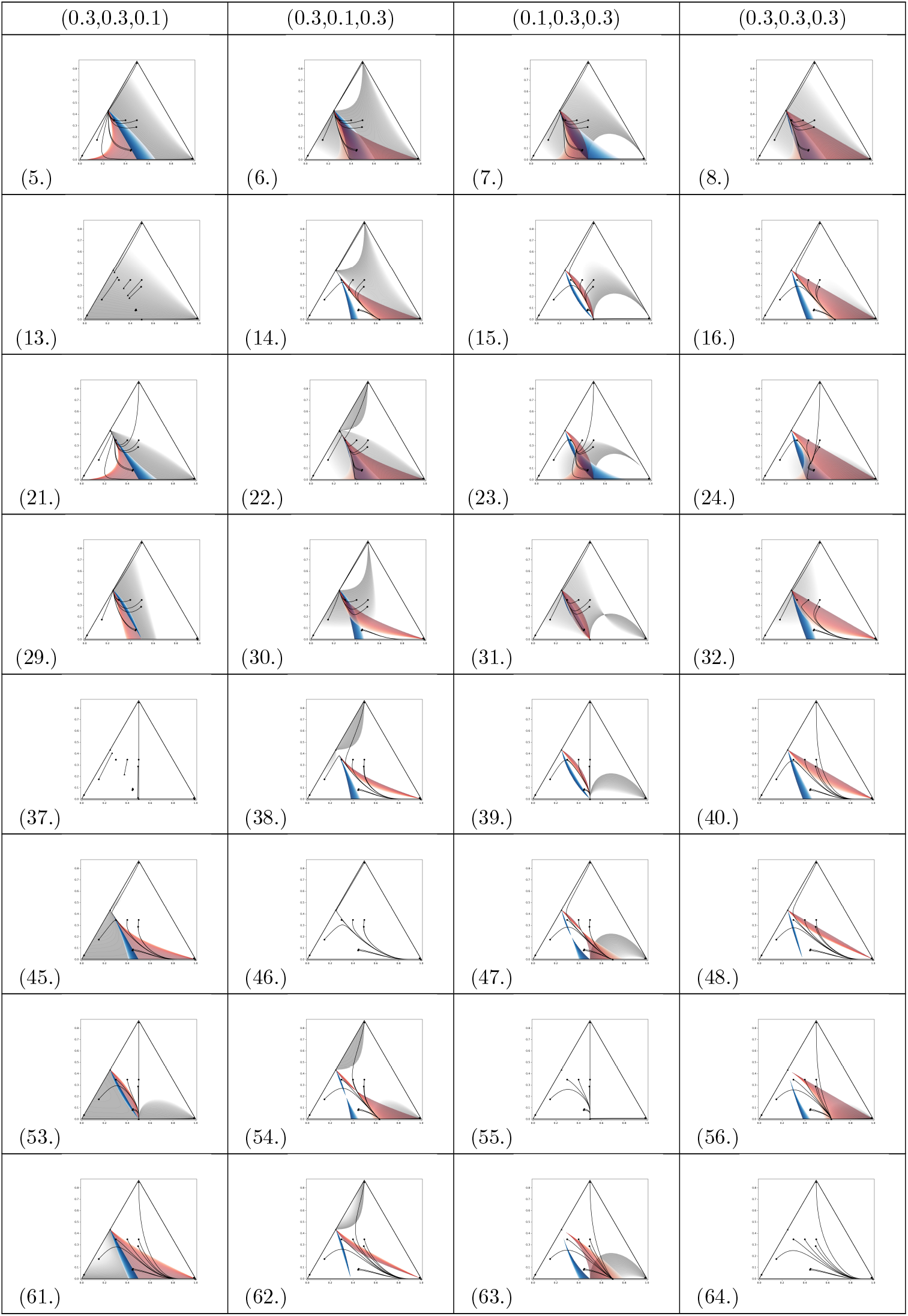

